# Diffusion of constructed resources and specialism promotes facilitation in spatial systems

**DOI:** 10.1101/2024.06.05.597587

**Authors:** Alice Nadia Ardichvili, Sébastien Barot, Jean-Christophe Lata, Nicolas Loeuille

## Abstract

Sessile organisms contend with locally limiting nutrients and neighbours. Increases in nutrient concentration due to niche construction and diffusion of resources change the interaction sign between neighbours usually thought to compete for resources. We investigate conditions under which the positive effect of niche construction outweighs the negative effect of resource consumption. Following the design of facilitation experiments, we model two patches connected by resource flows, and ask whether the presence of a niche constructor in one patch eases the colonisation of the other. Niche constructors positively affect neighbouring organisms when their niches are sufficiently differentiated, and when the constructed resource diffuses more than the non-constructed resource. Our work proposes mechanisms for the emergence of facilitation, which is increasingly recognised as a key process structuring plant and microbial communities. We discuss the implications for the spatial structure of communities and their functioning, including in an agricultural context.

## Introduction

Facilitation is widespread among plant species (Michalet and Pugnaire, 2016; Yin et al., 2022) and micro-organisms (Piccardi et al., 2019), potentially occurring at a local scale. The mechanisms driving facilitation are diverse and include the creation of a favourable microclimate (Callaway et al., 2002; Michalet et al., 2014), sediment retention (Bruno, 2000), the fixation of nitrogen (Koffel et al., 2018), and production of by-products used as resources in cross-feeding communities (Goldford et al., 2018). Interactions potentially switch from competition to facilitation as external stress increases (Callaway et al., 2002; Michalet et al., 2014; Piccardi et al., 2019), switch from facilitation to competition at different life stages (He et al., 2013; Yin et al., 2022), or at different stages of succession (Koffel et al., 2018). Because the nature of interactions ultimately drives the coexistence of species and the structure of ecological communities, understanding conditions for the emergence of facilitation remains an important stake for modern ecology.

Within communities, the effects of competition are often viewed in terms of niche differentiation in a local context where resources are shared (Chesson, 2000; Tilman, 1982). Competition is minimal when niches are sufficiently differentiated (stabilising mechanisms *sensu* Chesson, 2000). In ecosystems connected by resource flows, spatial heterogeneity changes the interaction sign between organisms, limiting the competition between organisms with similar niches and allowing their coexistence (Huston and DeAngelis, 1994), or limiting the exclusion of weaker competitors due to diffusion of resources (Gravel et al., 2010; Haegeman and Loreau, 2015). The dispersion modality of the consumers or the resources (informed, symmetrical) alters the interaction sign between organisms and ultimately the conditions of coexistence (Gravel et al., 2010; Haegeman and Loreau, 2015). These models highlighted the role of asymmetry in diffusion, but it is expected that niche construction, modifying resource concentration, also modifies the asymmetry in resource availability.

In the modern theory of coexistence (Barot and Gignoux, 2004; Chesson, 2000; Tilman, 1982), organisms are viewed as consumers depleting local resources; their ability to modify resource flows is ignored (but see Koffel et al., 2021). However, many niche-constructing activities increase the quantity of one resource (plants performing Biological Nitrogen Fixation, *Lupinus* species increasing the availability of phosphorus (Hinsinger et al., 2003), ants raising aphids for their nectar). Another assumption in the modern theory of coexistence is that resources (in particular their supply levels) are independent. Some nutrients are indeed completely independent and should be modelled as two independent resources (eg. phosphorus, potassium). However, some nutrients are taken up in several forms that are not independent when one form becomes the other (ammonium and nitrate as different sources of nitrogen for plants (Britto and Kronzucker, 2013), by-products as different sources of carbon for bacteria (Goldford et al., 2018)). In cases where one resource becomes the other, the possible effect of a niche constructor is to modify the rates of transformation, e.g. plant controlling nitrification (Lata et al., 2004), mineralisation rates (Shahzad et al., 2018). For example, the control of nitrification by plants changes the conditions of coexistence between species with distinct preferences for ammonium vs. nitrate (Ardichvili et al., 2024; Boudsocq et al., 2012).

With a theoretical approach, we determine the effect of a niche constructor influencing the dynamics of resources in its patch, on the neighbours. We seek the conditions in which a niche constructor has a facilitating effect on neighbours, i.e. eases their establishment during colonisation. We proceed by analogy with the experimental designs assessing facilitation, by testing whether the growth rate of a small population is positively or negatively impacted by its niche-constructing neighbour (Callaway et al., 2002). We restrict our analysis of niche construction on resources and do not take into account the modification of environmental conditions *sensu* (Chapin et al., 2002), such as the modification of micro-climate around plants. We vary the type of niche construction (accumulation of one resource, control of the transformation rate), the intensity of niche construction, the total flow of resources between patches (exchange rate), the propensity of one resource to diffuse more than the other (diffusion asymmetry), and the niche difference between the constructor and its neighbours.

We build a two-patch model, each containing two types of resources. In the first scenario, the resources are independent and the organisms simply consume the resources. In the second scenario, the niche constructor farms one resource and increases the rate of entry of that resource. In the third scenario, we consider two dependent resources, and the niche constructor modifies the rate of transformation of one resource to the next.

When an organism only consumes resources, the resource levels are necessarily lower in the occupied patch than in the empty patch, and the movement of resources is directed from the empty patch to the occupied patch, resulting in competition regardless of the niches of the two species. We expect that strong niche constructors invert this tendency and improve resource availability in the empty patch. However, the ability of a niche constructor to improve resource availability in the empty patch likely depends on its preference for the constructed resource and on the asymmetry in resource flow. We expect that the effect of the niche constructor switches from competition to facilitation when niche difference, niche construction, and diffusion of the constructed resource are sufficient.

## Methods

We model two patches containing two types of resources (Fig. 1).

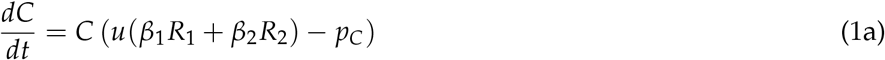

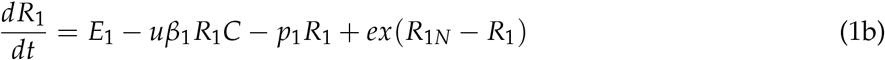

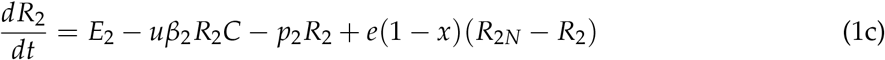

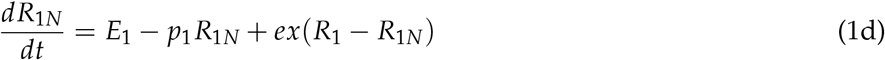

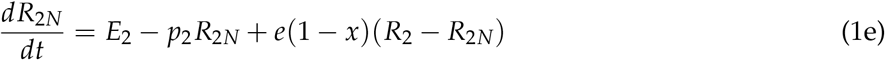

One patch is occupied by a niche constructor *C*, taking up resources *R*_1_ and *R*_2_ with a preference *β*_1_ and *β*_2_. We suppose that these resources are substitutable, thus *β*_1_ + *β*_2_ = 1. We consider continuous variations of *β*; when *β*_1_ ≈ 1, the organism is specialised on *R*_1_; when *β*_2_ ≈ 1, the organism is specialised on *R*_2_; and when *β*_1_ = *β*_2_ = 0.5, the organism has no preference and is a generalist. The neighbouring patch, initially unoccupied, also contains the same two resources *R*_1*N*_ and *R*_2*N*_ (Fig. 1.)

**Figure 1:**
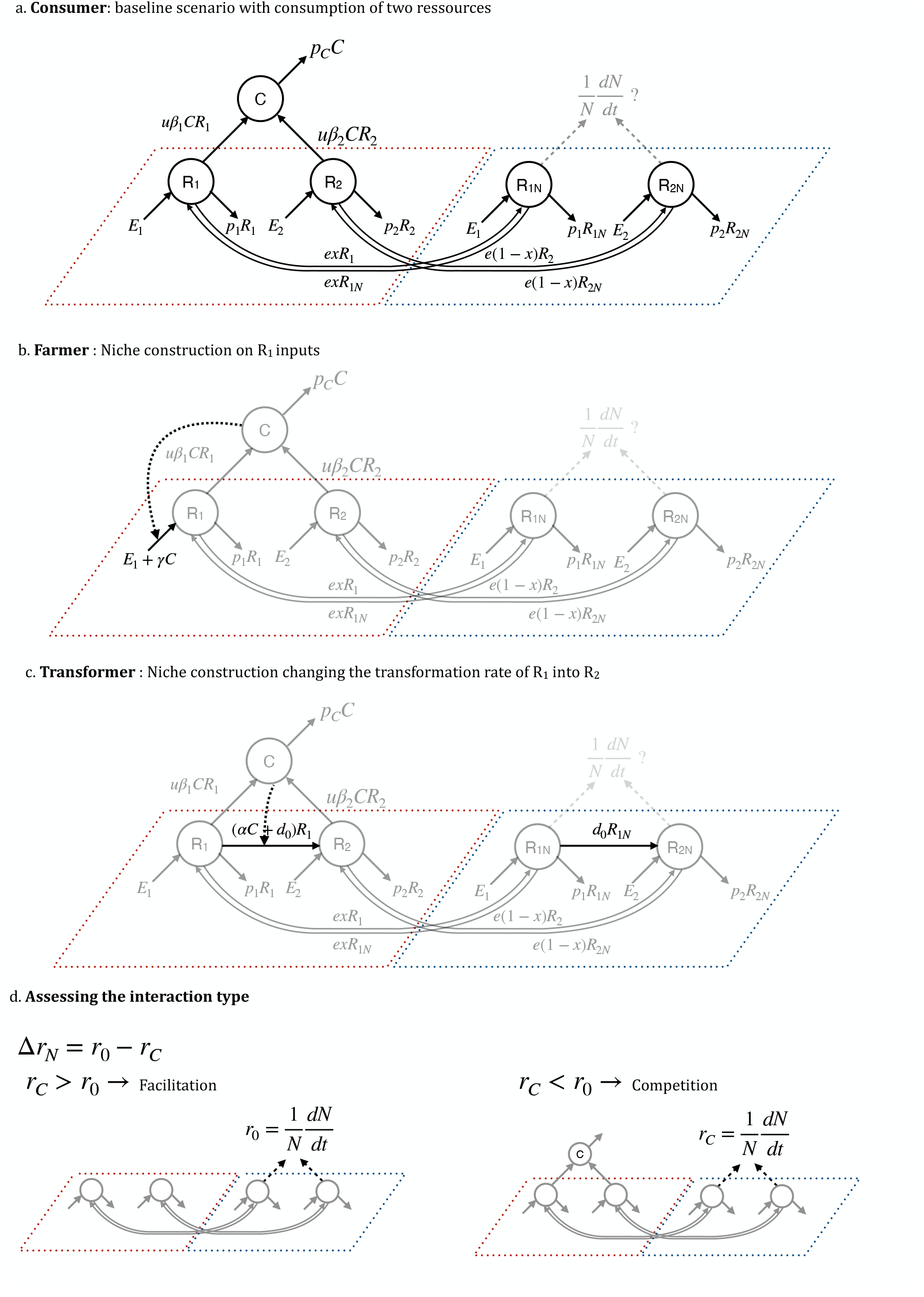
**Description of the model and the three niche construction scenarios. The niche constructor lives in the left patch (delimited by a red dashed line) and impacts, either positively or negatively, the growth of a coloniser** *N* **arriving in the neighbouring patch on the right (delimited by dashed blue lines). a. Baseline scenario: the niche constructor does not actively build its niche and consumes the two resources. b. The constructor increases the inputs of** *R*_1_ **at a rate** *γ*. **The flux that differs from the baseline model is in black, fluxes that are identical with the baseline scenario are in grey. c**. *R*_1_ **becomes** *R*_2_ **at a rate** *d*_0_, **and the constructor can modify this rate by its action (***α***). d. The methodology used to determine whether the niche constructor facilitates or hinders the neighbour is comparable to that of experimental designs (Callaway et al**., **2002); we conclude that the niche constructor has a facilitating effect when the growth rate of the coloniser is higher when the constructor is present compared to when the constructor is absent**.

We focus on the effects of niche construction, thus the two patches are identical (same input and loss rates in the resource compartments) and differ only by the presence of the niche constructor. The patches are connected by fluxes of resources flowing at a basal rate *e*, modulated by an asymmetry coefficient *x* varying continuously. When *x* = 1, only *R*_1_ diffuses; when *x* = 0, only *R*_2_ diffuses; when *x* = 0.5 the two resources diffuse at the same rate. Resources are taken up based on donor-controlled functions at a rate *u*. In the two patches, the resource dynamics are chemostat-like; the input rate is constant (*E*_*i*_) and losses proportional to the compartment size (*p*_*i*_ * *R*_*i*_, *p*_*C*_ * *C*, Fig. 1). Table 1 describes variables and parameters.

**Table 1:** Variables, parameters, units and domains.

|  | Variables | Unit | Domain |
| --- | --- | --- | --- |
| $t$ | time | yr | $\mathbb{R}^+$ |
| $C$ | biomass of the constructor organism | $\text{g m}^{-2}$ | $\mathbb{R}^+$ |
| $R_i$ | concentration of resource $i$ in the occupied patch | $\text{g m}^{-2}$ | $\mathbb{R}^+$ |
| $R_{iN}$ | concentration of resource $i$ in the neighbouring (and empty) patch | $\text{g m}^{-2}$ | $\mathbb{R}^+$ |
| Parameters |  |  |  |
| $u$ | uptake rate | $\text{g}^{-1} \text{m}^2 \text{yr}^{-1}$ | $\mathbb{R}^+$ |
| $E_i$ | input rate of resource $i$ | $\text{g yr}^{-1}$ | $\mathbb{R}^+$ |
| $p_i$ | loss rate of resource $i$ | $\text{yr}^{-1}$ | $\mathbb{R}^+$ |
| $p_c$ | loss rate of the constructor | $\text{yr}^{-1}$ | $\mathbb{R}^+$ |
| $\beta_i$ | preference for resource $i$ | - | $[0; 1]$ |
| $e$ | exchange rate between the patches (connectivity) | $\text{yr}^{-1}$ | $\mathbb{R}^+$ |
| $x$ | propensity of resource 1 to diffuse more than resource 2 | - | $[0; 1]$ |
| $\gamma$ | niche construction rate by the niche constructor | $\text{yr}^{-1}$ (Scenario 2) | $\mathbb{R}^+$ |
| $d_0$ | transformation rate of resource 1 to resource 2 (Scenario 3) | $\text{yr}^{-1}$ | $\mathbb{R}^+$ |
| $\alpha$ | control of the transformation rate of the constructor (Scenario 3) | $\text{g}^{-1} \text{m}^2 \text{yr}^{-1}$ | $\mathbb{R}$ such that<br>$C\alpha + d_0 > 0$ |

We use the baseline scenario, which does not contain niche construction, as a reference to study other scenarios.

In the “Farmer” scenario (eg. nitrogen fixation or mycorrhizal association, Fig. 1b), the niche constructor increases the input rate of *R*_1_ at a per capita rate *γ*, such that Eq.1b becomes:

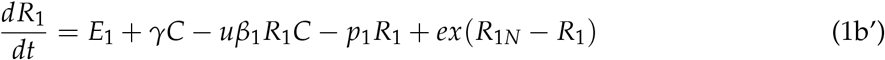

In the “Transformer” scenario (mineralisation of organic matter, control of nitrification rate, Fig. 1c), *R*_1_ becomes *R*_2_ at a basal rate *d*_0_. the niche constructor influences that rate; when *α* < 0, the organism inhibits the transformation (for example plants inhibiting nitrification); when *α* > 0, the constructor stimulates this transformation (for example plants boosting the mineralisation rate). Eq.1b-d become :

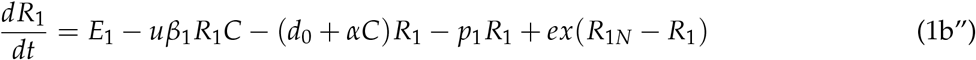

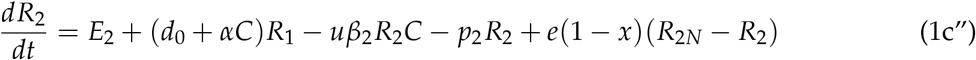

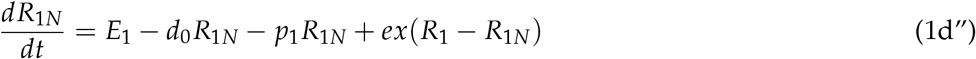

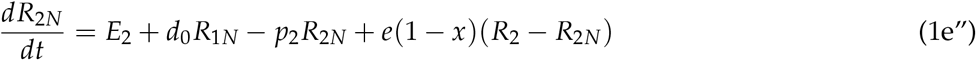

, keeping *d*_0_ + *αC* > 0 positive.

In this system, the growth rate of a neighbour arriving in the patch also depends on the uptake rate *u* and loss rate *p*_*C*_ (which are the same as the constructor’s), and the preferences for resources *β*_1*N*_ and *β*_2*N*_, and in the resource concentration *R*_1*N*_ and *R*_2*N*_ :

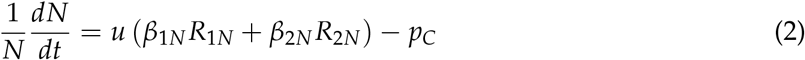

We consider several possibilities regarding the preferences of the neighbours, each allowing us to reach conclusions about different organisation scales. When *β*_1_ = *β*_1*N*_ and *β*_2_ = *β*_2*N*_, the two individuals are the same species and any phenotypic variability is ignored (population scale). When *β*_1_ ≠ *β*_1*N*_ and *β*_2_ ≠ *β*_2*N*_, the two individuals may be from the same species but the preference of one of them may have mutated (evolutionary scale). Alternatively, they may be individuals from ecologically close species, differing only in their preference, not in their uptake rate or mortality (community scale).

Determining the sign of the interaction between two species requires growing the species together and separately, either by removing neighbours in the field (Callaway et al., 2002) or by growing plants together and separately in pot experiments (Callaway, 2007). By analogy, to decide whether a niche constructor has a facilitating effect on its neighbour, we evaluate whether the growth rate of the neighbour is higher when the patch is occupied by a niche constructor (resource compartments stabilising at 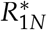 and 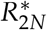 ) and when the neighbouring patch is empty (resource compartments stabilising at 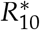 and 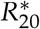), Fig. 1d.

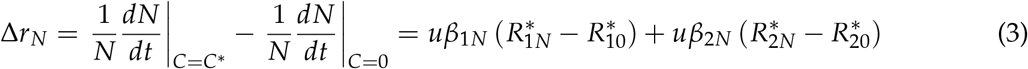

A constructor has a facilitating effect when the growth rate of the neighbour increases with the presence of the constructor, i.e. when Δ*r*_*N*_ is positive. We study the sign of Δ*r*_*N*_ and assess whether changes in interaction type depends on resource *R*_1_ our resource *R*_2_, ie. its components 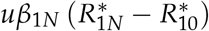 and 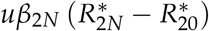.

We illustrate our findings using a graphical approach (Fig. 2) where we plot the concentrations of the two resources in the patches of the niche constructor and the coloniser. The concentration of the empty patch without diffusion (white circle) depends on the input and loss rates of the two resources. The concentration of the occupied patch, in the absence of diffusion, settles on the Zero Net Growth Isocline (ZNGI) of the niche constructor, i.e. the curve on which resource concentrations result in a null growth rate. We hypothesise that the type of niche construction influences resource concentration in the neighbour patch: the niche constructor either depresses the concentration of both resources and settles at the red circle, or increases the quantities of *R*_1_ (orange circle) or *R*_2_ (pink circle). Diffusion homogenises resource concentrations between the two patches, ie. bring the resource concentration of the empty patch (white circle) closer to that of the occupied patch (coloured circles). With diffusion, the concentration of the empty patch lands in the coloured rectangle, in a place that depends on the strength of diffusion (length of the coloured arrows) and on the asymmetry in diffusion (along the different arrows). The resulting concentrations either yield a higher growth rate for the coloniser (in the light green area of the plot) or a lower growth rate (light red area). The dashed grey line delimiting these two areas corresponds to the growth isocline sustaining the growth rate *r*_0_. Note that the steepness of this isocline and of the ZNGI of the niche constructor depend on their respective preferences (steeper growth isoclines for *R*_1_ specialists, flat growth isoclines for *R*_2_ specialists).

**Figure 2:**
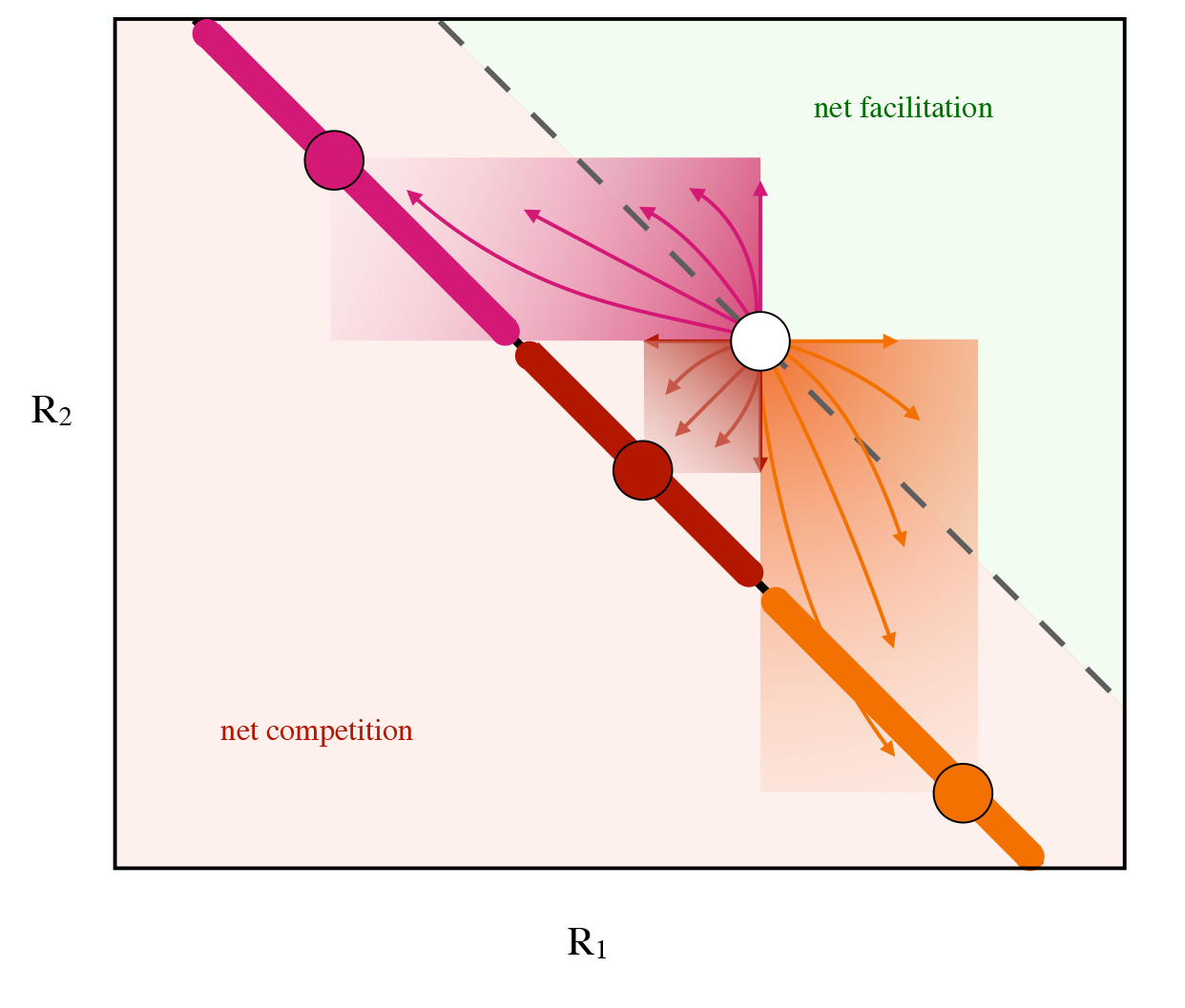
**Schematic representation of the conditions that allow facilitation. White circle: resource concentration in the empty patch without exchange or without a niche constructor. Dashed line: growth isocline of the neighbour at the positive growth rate** *r*_0_. **The angle of the isocline depends on the degree of specialism of the coloniser. Light green area: resource concentrations sustaining a growth rate larger than** *r*_0_**; light red area: resource concentrations resulting in a growth rate lower than** *r*_0_. **Thick coloured lines: ZNGI of the constructor. The three coloured circles illustrate the concentrations in the patches of three hypothetical niche constructors. The coloured squares indicate the possible concentrations that the neighbour patch takes as diffusion increases between the two patches. The coloured lines indicate hypothesised trajectories that the neighbour patch takes as diffusion increases (along the arrow) for different asymmetries (different arrows)**.

## Results

### Scenario 1: a consumer always negatively affects the neighbour

Studying the population of a niche constructor requires that the population exists, i.e. that a positive and reachable population equilibrium exists. The equilibrium is reachable when the population can invade an empty patch. The empty patch has concentrations of 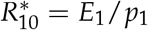 and 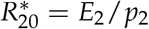. The growth rate of an initially rare niche constructor is positive when :

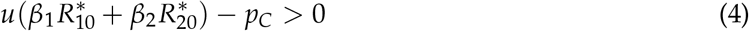

Eq. 4 means that the niche constructor exists when the capacity to acquire both resources is sufficient given its mortality. Then, only one equilibrium is feasible (see Appendix A).

At equilibrium, all derivatives are null and after manipulation (see Appendix B), we the effect of the consumer on the availability of the two resources for the neighbour as :

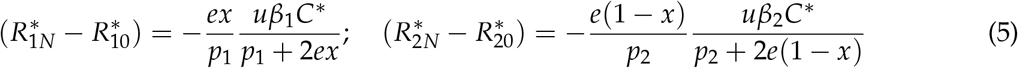

Since all parameters are positive, the two components of Δ*r*_*N*_ in Eq. 5 are negative. The consumer negatively impacts the neighbour but the magnitude of the negative effect depends on the preferences of the neighbour and the constructor, and the asymmetry in resource diffusion. When diffusion is completely asymmetric (*x* = 0 or *x* = 1), one of the terms becomes null, meaning that the strength of competition only depends on what happens with the diffused resource. If the neighbour is specialised on the non-diffused resource (*β*_1*N*_ = 1 and *x* = 0 or when *β*_1*N*_ = 0 and *x* = 1), the two terms become null, meaning that the constructor has a neutral effect on the neighbour. If the neighbour and the constructor are specialised on different resources (*β*_1_ = 0 and *β*_1*N*_ = 1 or *β*_1_ = 1 and *β*_1*N*_ = 0), the effect of the constructor is also neutral. Otherwise, the consumer is simply competing with the neighbour.

Figure 3a-b illustrates the negative effect of the consumer. The axes show the environmental state of the two resources (*R*_1_, *R*_2_) in the empty neighbouring patch in the absence of exchange (large white circle with coordinates 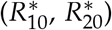) and in the patch occupied by the niche constructor (large black circle with coordinates 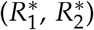). Here, the niche constructor is a simple consumer, so that the resource concentration is weaker in the occupied patch than in the empty patch (the black circle is below/left of the white circle). Consequently, diffusion lowers resources in the neighbour patch and the constructor harms the neighbour (competition, shown as red dots in Fig. 3). Every small circle, with coordinates 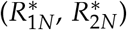 corresponds to the equilibrium state of the neighbouring patch when exchange is allowed as the exchange rate *e* increases, and are shown here for different asymmetries in diffusion. The colour of the point depends on the value of Δ*r*_*N*_, which is the effect of the constructor on the neighbour. Darker red circles indicate a stronger negative effect.

**Figure 3:**
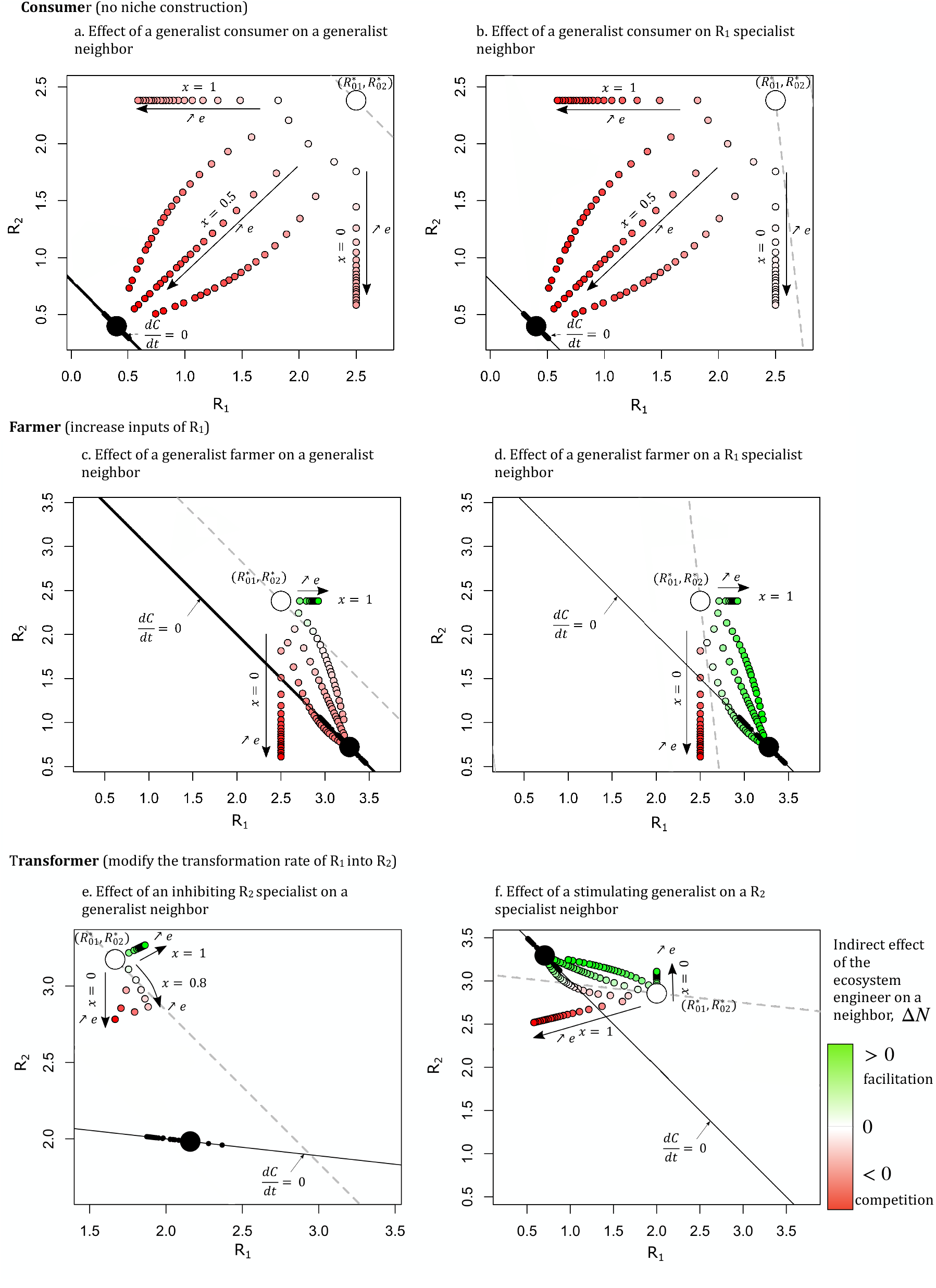
**The effect of the niche constructor depends on the type of niche construction, the magnitude** *e* **and asymmetry** *x* **of diffusion. Large circles: richness of the patch of the niche constructor (black) and of the neighbouring patch (white) at equilibrium, when there is no exchange between the patches. The black line is the Zero-Net-Growth Isocline of the niche constructor. The dashed grey line is the positive growth isocline of the neighbour such that the growth rate is the same as in the empty patch without exchange. Small circles: equilibrium richness of the neighbouring patch as exchange increases (from** 1 **to** 50 **along black arrows), for different diffusion asymmetries (along the different lines**, *x* = 0, 0.2, 0.5, 0.8 **or** 1**). The colour indicates whether in that situation the presence of the niche constructor is beneficial (**Δ*r*_*N*_ > 0, **green points) or detrimental (**Δ*r*_*N*_ < 0, **red points) to the establishment of the neighbour. a and b: in the ‘consumer’ scenario, the consumer always has a detrimental effect on the neighbour. c and d: a farmer (** *γ* = 1.8**) sometimes facilitates the establishment of the neighbour, especially when the neighbour is a** *R*_1_ **specialist (d). e and f: an inhibitor (***α* = −1, *d*_0_ = 1 **(e)) or a stimulator (***α* = 5, *d*_0_ = 0.5**) has a positive effect when the diffusion is biassed towards the resource whose density increases by the effect of niche construction. Parameters:** *E*_1_ = *E*_2_ = 5, *p*_*C*_ = *p*_1_ = *p*_2_ = 2, *u* = 5

Fig. 3a-b also illustrates the results of Eq. 3 and 5: increase in the exchange rate homogenises the resource concentrations between the consumer and neighbour patches, which is deleterious for the establishment of a neighbour. When the diffusion is completely asymmetric (*x* = 0 or *x* = 1), the concentration of one resource is not affected, and the circles are disposed along a vertical or horizontal line.

A specialist neighbour is more affected by the diffusion of its preferred resource. In Fig. 3a., the consumer and the neighbour are generalists and their growth isoclines are parallel. In Fig .3b., the consumer is a generalist (black line) while the neighbour is specialised on *R*_1_ (dashed grey line). The effect of the consumer is quantitatively different in both cases. When the neighbour is specialised on *R*_1_ and the diffusion is biassed towards *R*_2_ (small circles aligned vertically on panel a), the effect of the consumer is almost neutral (the circles are almost white), which is not the case when both the neighbour and the consumer are generalists (panel a).

### Scenario 2: a farmer possibly has a positive effect on its neighbour

In this scenario, the niche constructor increases the inputs of *R*_1_. The niche constructor reaches an equilibrium biomass, or grows exponentially depending on the intensity of niche construction (see Appendix A).

Whether the population reaches an equilibrium or grows exponentially, facilitation occurs due to the construction on *R*_1_, ie. 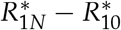 is positive, when (see Appendix B for derivation):

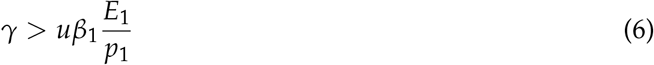

When condition (6) is fulfilled, farming leads to the accumulation of *R*_1_ in the local patch, fertilising the neighbour patch through diffusion. The niche construction rate *γ* needs to be sufficient compared to the capacity of the constructor to absorb the resource (*uβ*_1_) and the baseline richness of the empty patch (*E*_1_/*p*_1_). When *R*_1_ is not well conserved (*E*_1_/*p*_1_ is low), facilitation is likelier, which is in line with the *Stress Gradient Hypothesis* (Bertness and Callaway, 1994). When condition (6) is fulfilled, the large black circle in Fig. 3c-d. (state of the patch occupied by the constructor) is on the right side of the large white circle (state of the empty patch). When the constructor is specialised on *R*_1_, ie. high *β*_1_, the occupied patch is less rich than the empty patch both in terms of *R*_1_ and of *R*_2_, and the farmer behaves like a consumer of the previous scenario and has a negative effect (neutral at best) regardless of the diffusion and asymmetry. The effects of different preferences are explored visually in Appendix C.

To sum up, condition 6 shows that a niche constructor cannot facilitate the establishment of a neighbour, including a conspecific, when its preference for the constructed resource is too high. The rapid uptake of the farmed resource outweighs the constructing effect and the farmer only harms the neighbour. Facilitation occurs when the constructor sufficiently constructs *R*_1_ and does not have a strong preference for *R*_1_.

The effect of the niche constructor on *R*_2_, 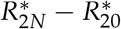, is the same as in Eq.5. There is always less *R*_2_ in the patch of the constructor due to the consumption by the constructor. The resulting competitive effect is shown in Fig. 3c-d. where the large black circle (state of the patch occupied by the constructor) is always below the large white circle (state of the empty patch). Therefore, in this scenario, the net interaction depends on the relative weight of the *R*_1_ term (facilitative when (6) is fulfilled) and the *R*_2_ term (always linked to higher competition).

When condition 6 is fulfilled, the exchange rate and asymmetry in diffusion determine whether facilitation occurs (Fig. 3). As in the consumer scenario, exchange between patches homogenises resource concentrations in a way that depends on asymmetry. Increase of the exchange rate is beneficial for the neighbour (small green circles) as long as the resulting resource concentration in the neighbour patch is above the dashed grey line. When the constructor and the neighbour have the same niche (Fig. 3c.), the effect of the constructor is positive only when the exchange is strongly biassed towards the constructed resource *R*_1_ (*x* = 1 at the horizontal green line, or *x* = 0.8 at the two pale green circles). When the diffusion is not 100% asymmetric, the exchange rate has to be sufficiently weak as to limit losses of *R*_2_ (the two circles are the closest to the large white circle and correspond to weak exchange rate *e*). To summarise, a constructor has a positive effect, including on a conspecific neighbour, when the resource that it farms is the resource diffusing the most.

When the neighbour is specialised on the constructed resource *R*_1_ (Fig. 3d.), more combinations of exchange and asymmetry are beneficial for the neighbour. In panel d, the farmer has a negative effect when *R*_2_ only diffuses (vertical line of red circles at *x* = 0). On the contrary, when the neighbour has a stronger preference for *R*_2_ (Fig.C2f and k), facilitation is less frequent. Facilitation by the constructor is possible for a larger range of exchange and asymmetry when species or other phenotypes are specialised on the farmed resource.

Appendix D explores the combinations of exchange rate *e* and asymmetry *x* leading to facilitation by the constructor. The more the diffusion is biassed towards *R*_1_, the more the presence of the constructor is beneficial for the neighbour. The effect of the exchange rate is more ambiguous and interacts with asymmetry (Fig. D1). When the farmed resource diffuses the most, the constructor has a positive effect on the neighbour for low exchange rates. On the contrary, when the non-farmed resource diffuses most, the exchange rate needs to be sufficiently large for facilitation to happen. When the neighbour is a specialist of the farmed resource (Fig. D1b), increase in the exchange rate *e* is always beneficial.

### Scenario 3: inhibitors can facilitate R_2_ specialists

In the last scenario, we consider a niche constructor influencing the transformation of *R*_1_ into *R*_2_ (Eq. 1b”-1e”), either by inhibiting the transformation (*α* < 0) or by stimulating it (*α* > 0). The population reaches a positive equilibrium, grows exponentially or exhibits bi-stable dynamics (see Appendix A for more details).

When the population reaches an equilibrium, the first component of Δ*r*_*N*_ reads :

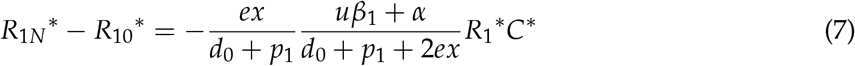

A stimulator (*α* > 0) has naturally a negative effect on the richness of *R*_1_ in the neighbouring patch due to its consumption of the resource and its transforming effect. As for an inhibitor (*α* < 0), the intensity of niche construction needs to outweigh the consumption of *R*_1_ (such that −*α* > *uβ*_1_) for the first component of Δ*r*_*N*_ to be positive.

When the population reaches an equilibrium, the second component of Δ*r*_*N*_ reads :

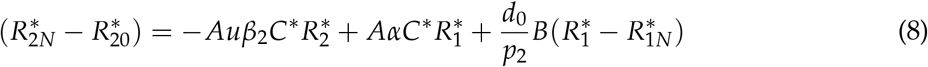

with

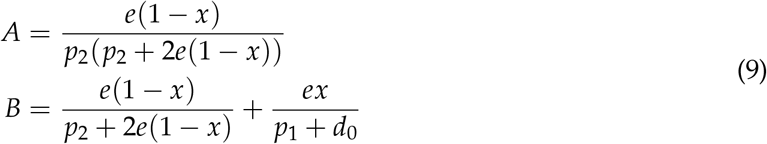

*A* and *B* are both positive. The first term, 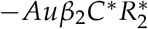 is the negative effect of the constructor due to the consumption of *R*_2_. The second term, 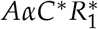 corresponds to the direct effect of the constructor on *R*_2_ due to its niche construction activity; it is positive when the constructor is a stimulator and negative when the constructor is an inhibitor. The last term, 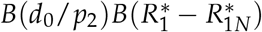, corresponds to the indirect niche construction effect via the modification of *R*_1_. This term is always negative when the niche constructor is a stimulator but can be positive when the niche constructor is an inhibitor. In other words, an inhibitor potentially increases the concentration of *R*_2_ in the empty patch. It is especially the case when diffusion is biassed towards *R*_1_: the term *A* becomes negligible and only the positive term multiplied by *B* remains. This positive indirect effect is illustrated in Fig. 3e. where, when x=1, the concentration of the neighbour patch increases in *R*_1_ and *R*_2_ content (small green circles moving up and to the right as exchange increases). Reciprocally, a stimulator negatively affects *R*_2_ due to its indirect effect on *R*_1_.

Fig. 3e shows that as in the “Farmer” scenario, an inhibitor has a positive effect on the neighbour when the diffusion is biassed towards *R*_1_. Symmetrically, a stimulator has a positive effect when diffusion is sufficiently biassed towards *R*_2_ (Fig 3f). Due to the transformation of resources, as opposed to the first two scenarios, diffusion of *R*_1_ only changes the patch concentrations of both *R*_1_ and *R*_2_ (circles at *x* = 1 are not horizontal in Fig. 3e and f).

As in the farmer scenario, for a given asymmetry in diffusion *x*, an increase in the exchange rate is beneficial or harmful to the establishing neighbour. In Appendix C3h, for *x* = 0.8, the presence of the transformer is beneficial at weak diffusion and becomes detrimental when *e* increases. In Appendix C4f, for *x* = 2, the constructor has a detrimental effect on the coloniser for a weak exchange rate and is beneficial when the exchange rate increases. The effect of the exchange rate depends on the asymmetry in resource diffusion.

As in the “Farmer” scenario, whether facilitation occurs depends on the degree of specialism of the two species. In most combinations explored in Appendix C, (Fig. C3 for an inhibitor and C4 for a stimulator), the niche constructor has a positive effect if niches are sufficiently differentiated, and if the coloniser is specialised on the constructed resource (*R*_1_ for an inhibitor, *R*_2_ for a stimulator). A notable exception is the case of two *R*_2_ specialists when the constructor is an inhibitor: facilitation occurs when *R*_1_ diffuses only (Fig. C3i). Although the two species have the same preferences, facilitation is possible because the inhibitor increases *R*_1_, and ultimately *R*_2_ concentrations after its transformation in the empty patch.

## Discussion

Facilitation and competition both structure plant and microbial communities (D’Andrea et al., 2024; Yin et al., 2022). Our work aims to clarify the conditions in which the consumption of resources and niche construction result in facilitation or competition. We used a two-patch model to study whether a niche constructor facilitates the establishment of a neighbour *via* the modification of resource availability, and under which conditions (Table. 2). We illustrate our findings using a graphical approach. According to our definition of facilitation, facilitation occurs when resource concentration in the neighbour patch is higher in the presence of a constructor. Diffusion homogenises the concentrations between the occupied patch and the neighbour patch. The concentration of the latter lands on the ZNGI of the niche constructor. The concentration of the neighbour patch naturally lands in the coloured rectangles depending on the exchange rate and the asymmetry in resource diffusion. A cultivator, increasing the input rate of one resource, potentially has a facilitating effect when niche construction is sufficient and basal resource availability is low. On top of these two necessary conditions, the exchange rate and diffusion asymmetry parameters need to fall in a place where the quantity of constructed resources brought in the empty patch outweigh the quantity of non-constructed resource (i.e. the resource that is consumed only) leaving the patch due to diffusion. This happens when asymmetry is strongly biassed towards the constructed resource, or when the exchange rate is weak and only slightly biassed towards the constructed resource. The conditions of facilitation in a situation where one resource becomes another are qualitatively similar; when the niche constructor increases the rate of transformation of one resource into another, the constructors acts like a farmer increasing the quantity of the second resource; when the niche constructor decreases the rate of transformation, the constructor acts like a farmer increasing the concentration of the first resource.

In line with a general theory of the niche (Koffel et al., 2021), our results show that the preferences of the neighbour and the niche constructor determine whether the constructor has a facilitating effect. Neighbour specialism on the constructed resource and specialism on the non-constructed resource on behalf of the niche constructor increases the possibilities of facilitation. When the organisms have the same preferences (because they belong to the same species, for example; panels a, e and i in Appendix C), facilitation is possible in rare cases: when both organisms are specialised on the non-constructed resource and when diffusion is strongly biassed towards the constructed resource. For example, nitrate in the soil is much more mobile than ammonium. The stimulation of nitrification, performed for example by trees in west-African savannas (Srikanthasamy et al., 2018) could result in intra-specific facilitation. The spatial aggregation of nitrification-stimulating trees further enforces the idea that facilitation occurs between such trees.

**Table 2:**
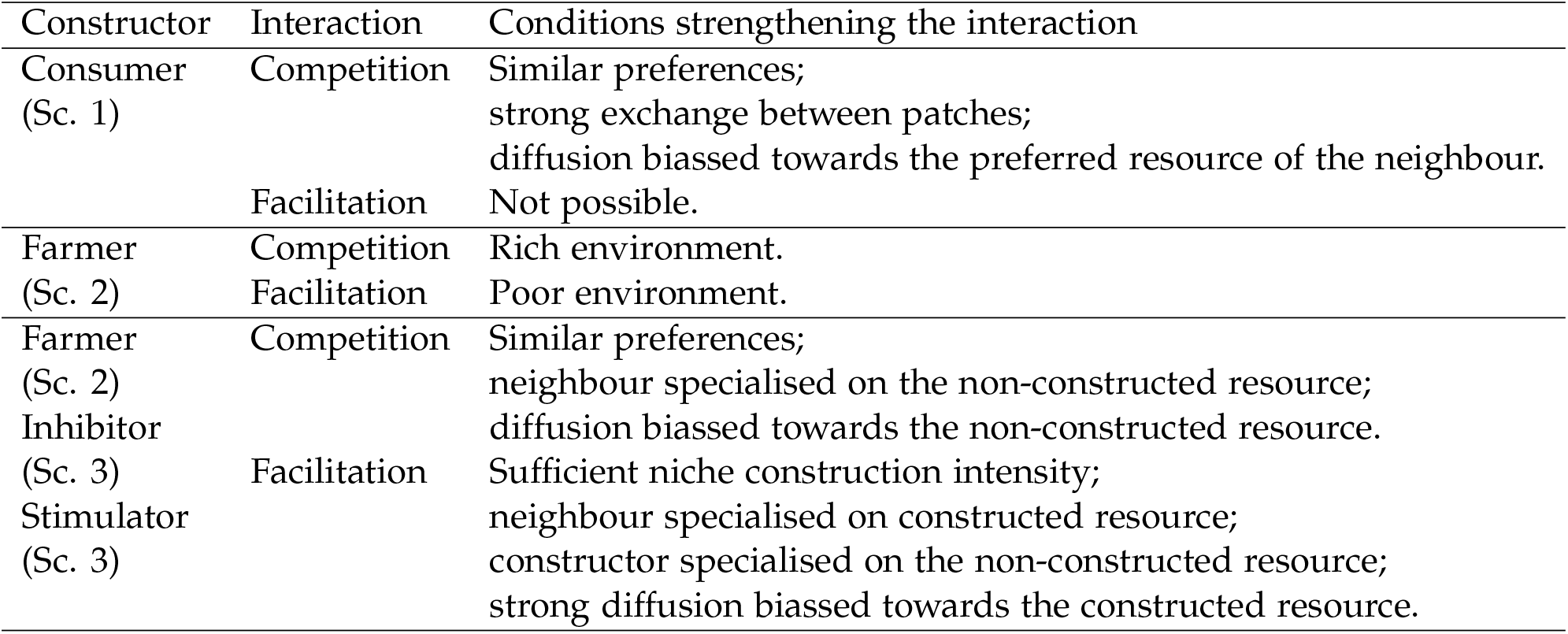
Summary of results: conditions that favour competition and facilitation in the three scenario.

| Constructor | Interaction | Conditions strengthening the interaction |
| --- | --- | --- |
| Consumer<br>(Sc. 1) | Competition | Similar preferences;<br>strong exchange between patches;<br>diffusion biased towards the preferred resource of the neighbour. |
|  | Facilitation | Not possible. |
| Farmer<br>(Sc. 2) | Competition | Rich environment. |
|  | Facilitation | Poor environment. |
| Farmer<br>(Sc. 2)<br>Inhibitor<br>(Sc. 3)<br>Stimulator<br>(Sc. 3) | Competition | Similar preferences;<br>neighbour specialised on the non-constructed resource;<br>diffusion biased towards the non-constructed resource. |
|  | Facilitation | Sufficient niche construction intensity;<br>neighbour specialised on constructed resource;<br>constructor specialised on the non-constructed resource;<br>strong diffusion biased towards the constructed resource. |

When the two organisms have different preferences (for example when individuals are of different species), a niche constructor has a facilitating effect when the constructed resource is more beneficial to the neighbour (of another species, ie. with a different preference) than to the constructor. A prerequisite of facilitation is that the niche of the neighbour and the constructor are sufficiently differentiated. A larger niche sensitivity difference *sensu* Koffel et al. (2021) increases possibilities of mutualism in a general model (Koffel et al., 2021). Tree-grass coexistence may also increase due to tree (grass) stimulating (inhibiting) nitrification in a spatial model of West-African savannas (Konare et al., 2024). During the first steps of ecological succession, phosphorus-limited species fixing nitrogen facilitate the establishment of species that are less limited by phosphorus but require more nitrogen (Bytnerowicz and Menge, 2021; Koffel et al., 2018).

Supposing that preferences are slightly different, as is the case in an evolving population where the neighbour has mutated, the constructor facilitates neighbours having a stronger preference for the constructed resource. In an adaptive dynamics framework (Dieckmann and Law, 1996), the successful colonisation of such mutants may result in the replacement of the resident population and lead to the directional selection of preference. In our model, the niche constructors would become specialised on the constructed resource in the long term. The joint evolution of preference and niche construction resulting in specialisation on constructed resources has also been observed in theoretical studies (Krakauer et al., 2009; Picot et al., 2021) and in the empirical system of farming ants becoming dependent on their farmed mushroom (Cafaro and Currie, 2005). In a microbial system, the co-evolution of two species leads to the specialization on the resource that is constructed by the other species resulting in mutualism between species (Harcombe, 2010).

We found that some conditions for facilitation are in line with the Stress Gradient Hypothesis (Bertness and Callaway, 1994; Callaway et al., 2002). In the second scenario, in which the constructor increases the entry of one resource, facilitation is more likely when the ecosystem is poor, i.e. more stressful (*E*_1_/*p*_1_ is weak). The SGH has been empirically derived from the observation of plants that create a favourable microclimate in cold environments. Our findings provide theoretical justification for the SGH in resource gradients which has been documented in experimental systems (Maestre et al., 2009; Piccardi et al., 2019; Rebele, 2000).

Unlike previous models of niche construction (Krakauer et al., 2009; Kylafis and Loreau, 2011), we did not assume that niche construction is costly for the constructor. However, some niche-constructing mechanisms may be costly (Bouma et al., 2005; Kuzyakov and Domanski, 2000). We could model a cost by reducing the uptake rate *u* or by increasing mortality *l*_*C*_ of the niche constructor in proportion to the investment in niche construction. Increasing mortality or reducing the uptake rate would shift the ZNGI of the constructor up/right because the constructor would need more resources to compensate for the cost of niche construction. At equilibrium, the resource concentration of the constructor would increase, implying that the difference between the richness of the empty patch and of the occupied patch would decrease. Modelling a cost would not change the qualitative results but would mitigate the effects (positive or negative) of the constructor on the neighbour, and result in more neutral interactions.

Note that we model a particular type of niche construction that only manipulates resource availability. Niche construction includes multiple processes, including those that modify environmental conditions that are not resources (eg. that are not consumed such as micro-climate, pH…) (Odling-Smee et al., 1996). While our work does not explicitly model such type of niche construction, the modification of environmental conditions such as temperature, pH, or humidity could potentially affect the activity of decomposers and impact resource fluxes.

Conditions allowing local facilitation could influence the emergence of spatial patterns. In spatially explicit models of niche construction (Dong, 2022; Klausmeier, 1999; Lejeune et al., 2002; Rietkerk et al., 2002; Scanlon et al., 2007), the level of niche construction by niche constructors determines the spatial patchiness of organisms. In another study, the level of facilitation determines the spatial aggregation of organisms (Kéfi et al., 2007). Our study makes the direct link between niche construction and facilitation at a local scale, and therefore suggests that spatial patterns may emerge when species modify resource availability.

To test the prediction of the model, we could combine data on the identity of plant species facilitating others (for example from Yin et al. (2022)) with plant traits (whether they engage in symbiotic fixation, mycorrhizal associations, preference for different types of resources). We could therefore check that niche constructors have a facilitating effect only when their preference for the constructed resource is sufficiently weak. Experimentally, we could test the effect of diffusion using transplantation, as in plant-soil feedback experiments (van der Putten et al., 2013), and control the homogenisation between two pots by mixing different fractions of soil in each treatment.

From an agronomic perspective, knowing the conditions in which niche constructors have a positive effect on their neighbour could guide the choice of companion planting, cover crop plants or plants to be used in intercropping. To reduce synthetic inputs and improve nutrient cycling in agricultural landscapes, agroecologists suggest using companion plants, i.e. growing a target plant with a neighbour of a different species improving the growth conditions of the target plant (e.g. repelling predators, improving nutrient availability, Wezel et al. (2020)). Examples include growing cereals with N-fixing legumes (Layek et al., 2018), legumes with vegetables (Brooker et al., 2015), maize with *Braccharia*, a genera inhibiting nitrification (Souza et al., 2024). Our results suggest that the niches of the companion plant and the target plant, as well as the mobility of the resource mobilised by the companion, need to be taken into account to evaluate whether the companion effectively improves the yield of the target plant.

## Acknowledgments

We thank Thomas Koffel, Vincent Calcagno and Sonia Kefi for interesting discussions on the model.

## Appendix A: feasibility analysis in the three scenario

From the model equations (1, 1b’ and (1b”-1e”), for every model the population equilibrium *C*^*^ is the solution of a polynomial :

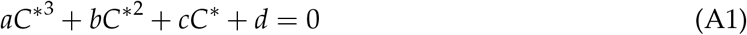

We use Descartes’ rule to find a positive and real equilibrium. In the three scenario,

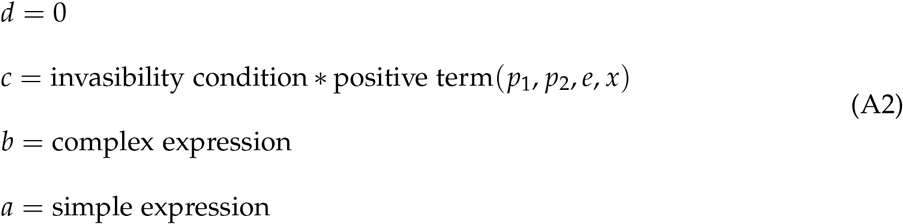

*c* is always positive when the invasibility condition is met.

Studying the sign of *a* allows us to determine when an equilibrium is feasible.

In the three scenarios, *a* reads *a*_*conso*_, *a*_*γ*_, and *a*_*α*_ for the consumer, farmer and transformer scenarios respectively.

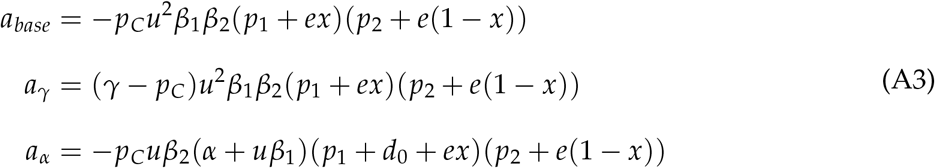

*a*_*conso*_ is always negative, *a*_*γ*_ is negative when *p*_*C*_ > *γ, a*_*α*_ is negative when *α* + *uβ*_1_ > 0. The sign of *b* does not inform us about the issue of the population dynamics since Descartes’s rule cannot say whether there are - or 2 positive roots when there are two sign changes. The existence of 1, 2 or 0 equilibria (number in red) results in the following dynamics (in black):

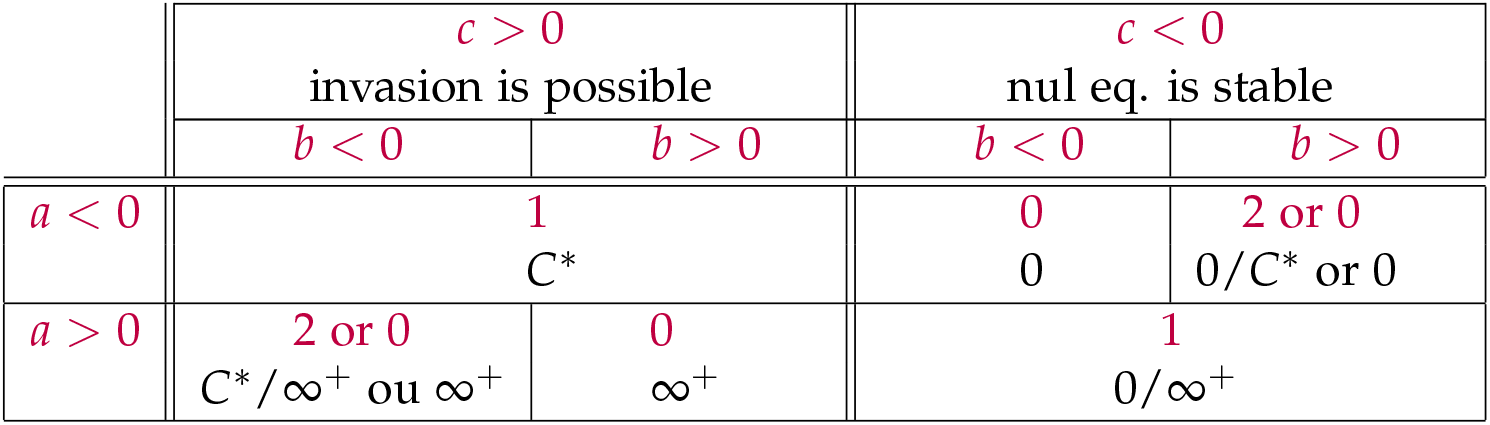

*C*^*^ indicates that the population stabilises at an equilibrium, 0 that extinction is locally stable, ∞^+^ the the population grows exponentially. The / indicates situations of bi-stability where the dynamics depend on the initial abundance of the niche constructor.

## Appendix B: obtaining equations 5 and 6

*Equation 5*

At equilibrium, *R*_10_ = *E*_1_/*p*_1_, Eq. 1d = 0 and:

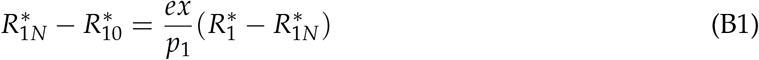

Assuming equilibrium, subtracting Eq. 1d to Eq. 1b yields

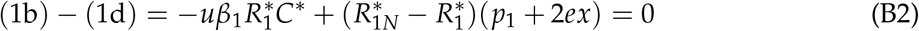

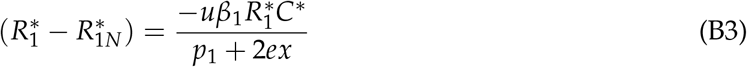

Subbing Eq.B3 into Eq. B1 yields Eq. 5.

*Equation 6*

In the second model, assuming equilibrium Eq. 1b’ and 1d can be re-arranged:

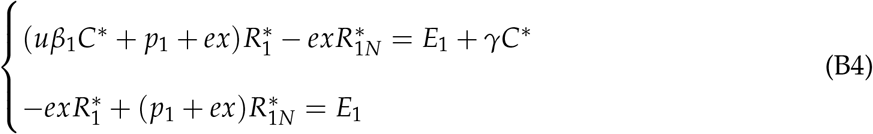

In matrix form, that yields

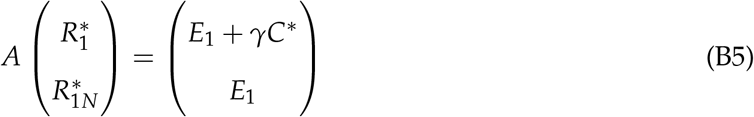

with

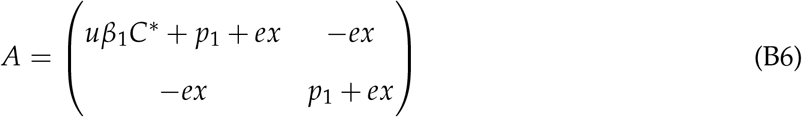

And

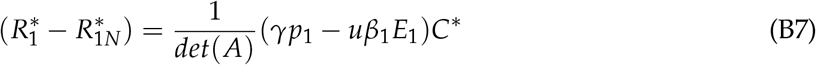

Where *det*(*A*) = (*p*_1_ + *ex*)*C*^*^*uβ*_1_ + *p*_1_(*p*_1_ + 2*ex*) is positive, so from Eq. B7 and B1 yield condition 6 when the niche constructor reaches an equilibrium (*C*^*^ > 0).

When the niche constructor grows exponentially,

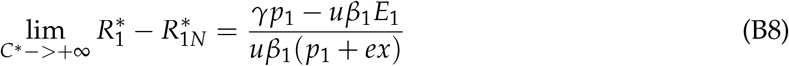

and the same condition holds.

*Equation 7*

At equilibrium, *R*_10_ = *E*_1_/(*p*_1_ + *d*_0_), Eq. d” = 0 and:

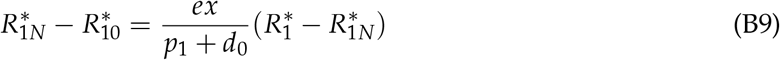

Subtracting 1d” to 1b” :

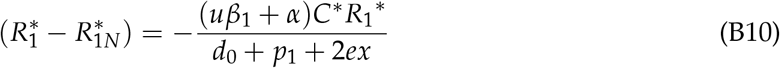

*Equation 8*

At equilibrium, 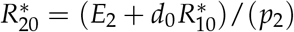.

Equation 1e” can be written :

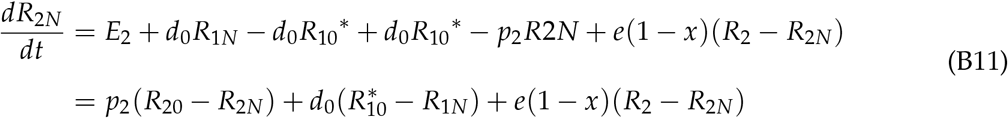

At equilibrium, Eq. B11 = 0 and we can write:

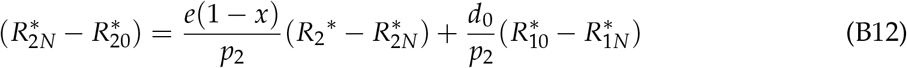

Subtracting 1e” to 1c” and assuming equilibrium, we obtain:

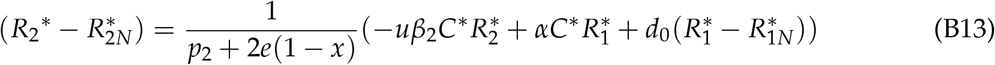

After re-arranging the previous 4 equations, we get :

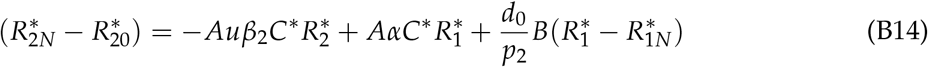

with

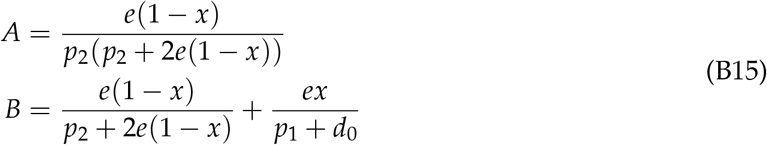

*A* and *B* are both positive. The first term, 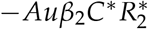 is the negative effect of the constructor due to the consumption of *R*_2_. The second term, 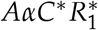 corresponds to the direct effect of the constructor on *R*_2_ due to its niche construction activity; it is positive when the constructor is a stimulator and negative when the constructor is an inhibitor. The last term, 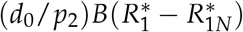, is always negative when the niche constructor is a stimulator but can be positive when the niche constructor is an inhibitor. In other words, even an inhibitor can increase the concentration of *R*_2_ in the empty patch.

## Appendix C: Effect of the niche constructor for different neighbour and constructor preferences

This appendix explores the effect of the niche constructor for different preferences *β*_1_ and *β*_1*N*_. Figure 3 in the main text shows several cases of facilitation. However, looking in more detail into several combinations of strategies highlight that cases of facilitation are rare.

First, facilitation is impossible when the constructor is specialised on the resource whose concentration increases due to niche construction (*R*_1_ in the Farmer scenario or when the Transformer is an inhibitor; *R*_2_ when the transformer is a Stimulator), which is in line with Eq. 6 and Eq.7. In other cases, facilitation is only possible when diffusion is sufficiently biassed towards the constructed resource. It’s only when the constructor and the neighbour have sufficiently different niches that a broader range of exchange and asymmetry allow facilitation.

**Figure C1:**
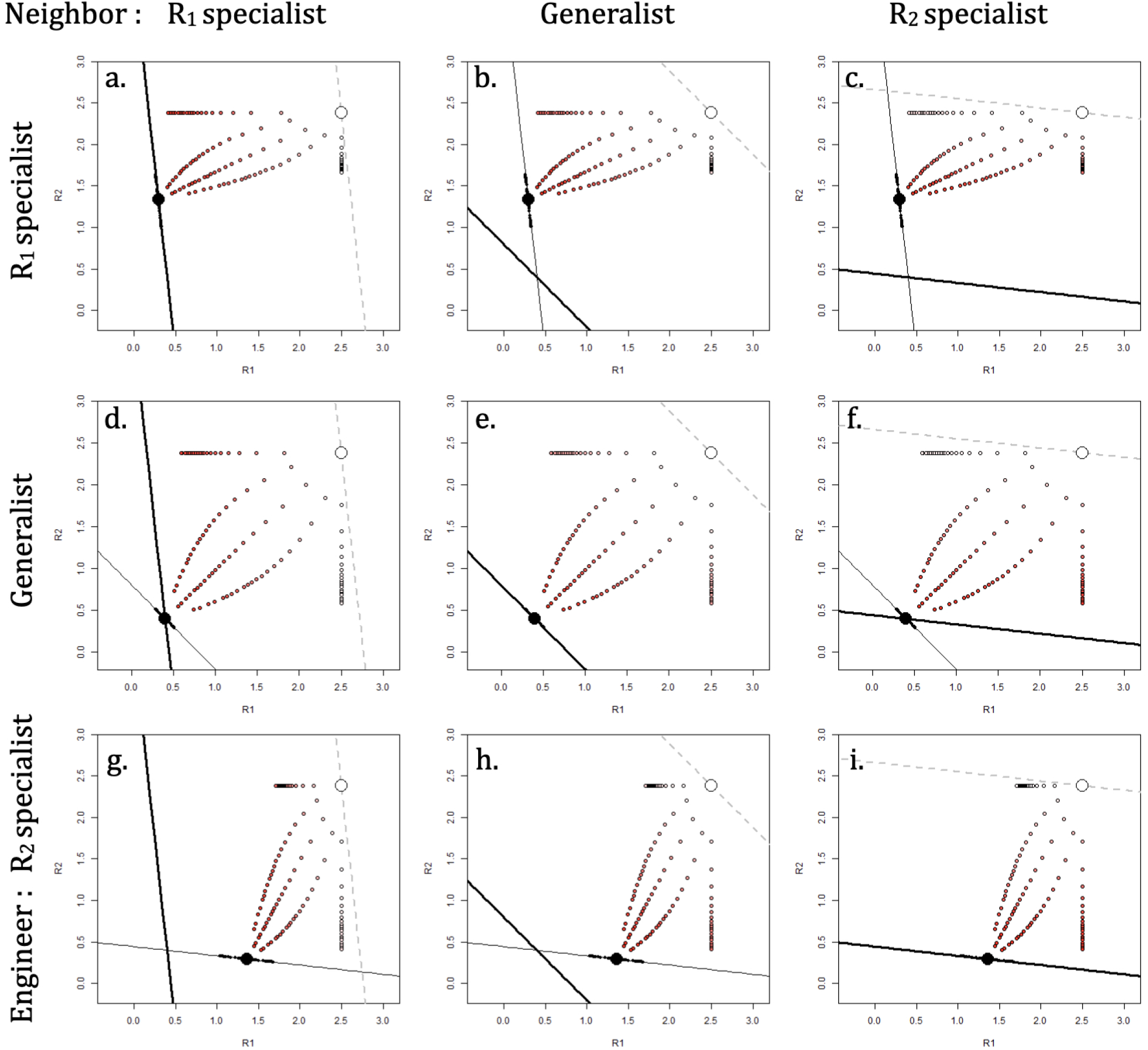
**Effect of the exchange rate and diffusion on the richness of the neighbour patch in the base model**. Δ*r*_*N*_ **is negative in red. As expected, whatever the strategies of the constructor or the neighbour, the effect is always negative. The colour scale is panel-dependent, darkest red corresponding to the worst case for the given** *β*_1_ **and** *β*_1*N*_ **combination**.

**Figure C2:**
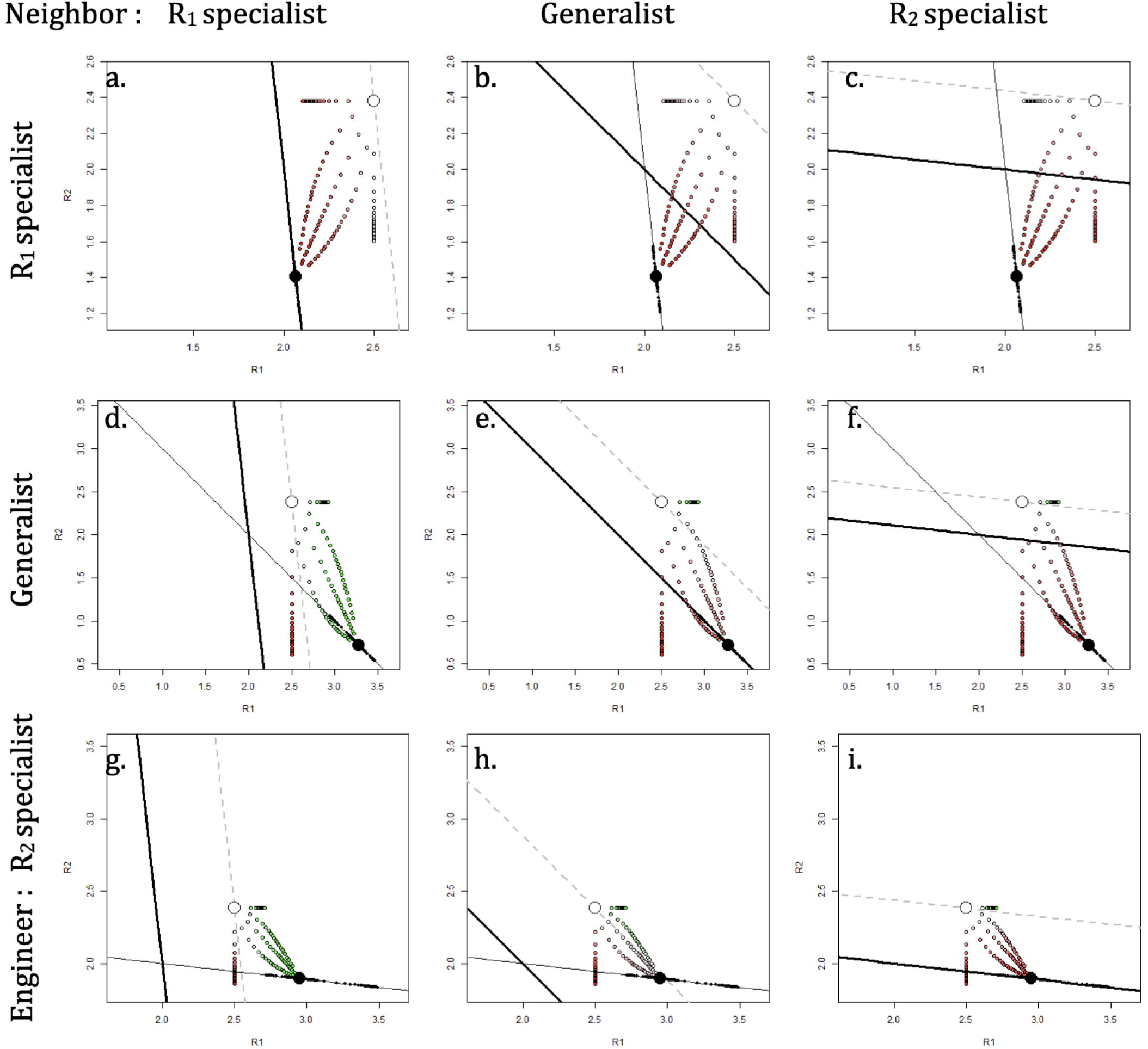
**Effect of a niche constructor farming resource 1**. Δ*r*_*N*_ **positive when the circle is green, negative when the circle is red. When the constructor is specialised on resource 1 (1st line), facilitation cannot happen (see condition 6). Facilitation is more likely when the neighbour has a strong preference for resource 1 (panels d and i). Level of facilitation :** *γ* = 1.8

**Figure C3:**
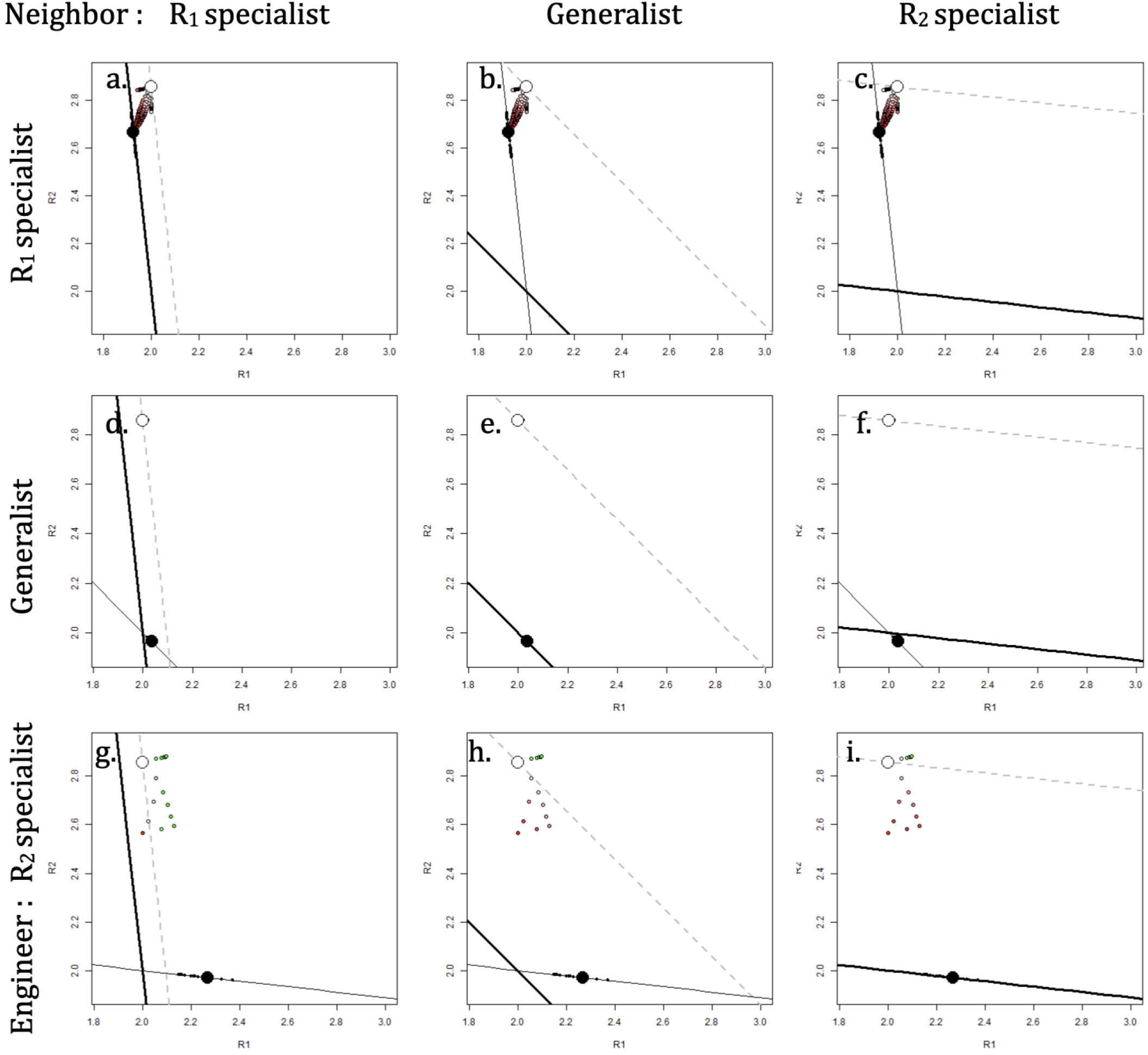
**Effect of an constructor slowing down the transformation of** *R*_1_ **into** *R*_2_. **As in the Farmer scenario, facilitation occurs only when the constructor is not a** *R*_1_ **specialist. Additionally, the neighbour needs to be a** *R*_1_ **specialist and only** *R*_1_ **should diffuse**. *d*_0_ = 0.5, *α* = −0.55

**Figure C4:**
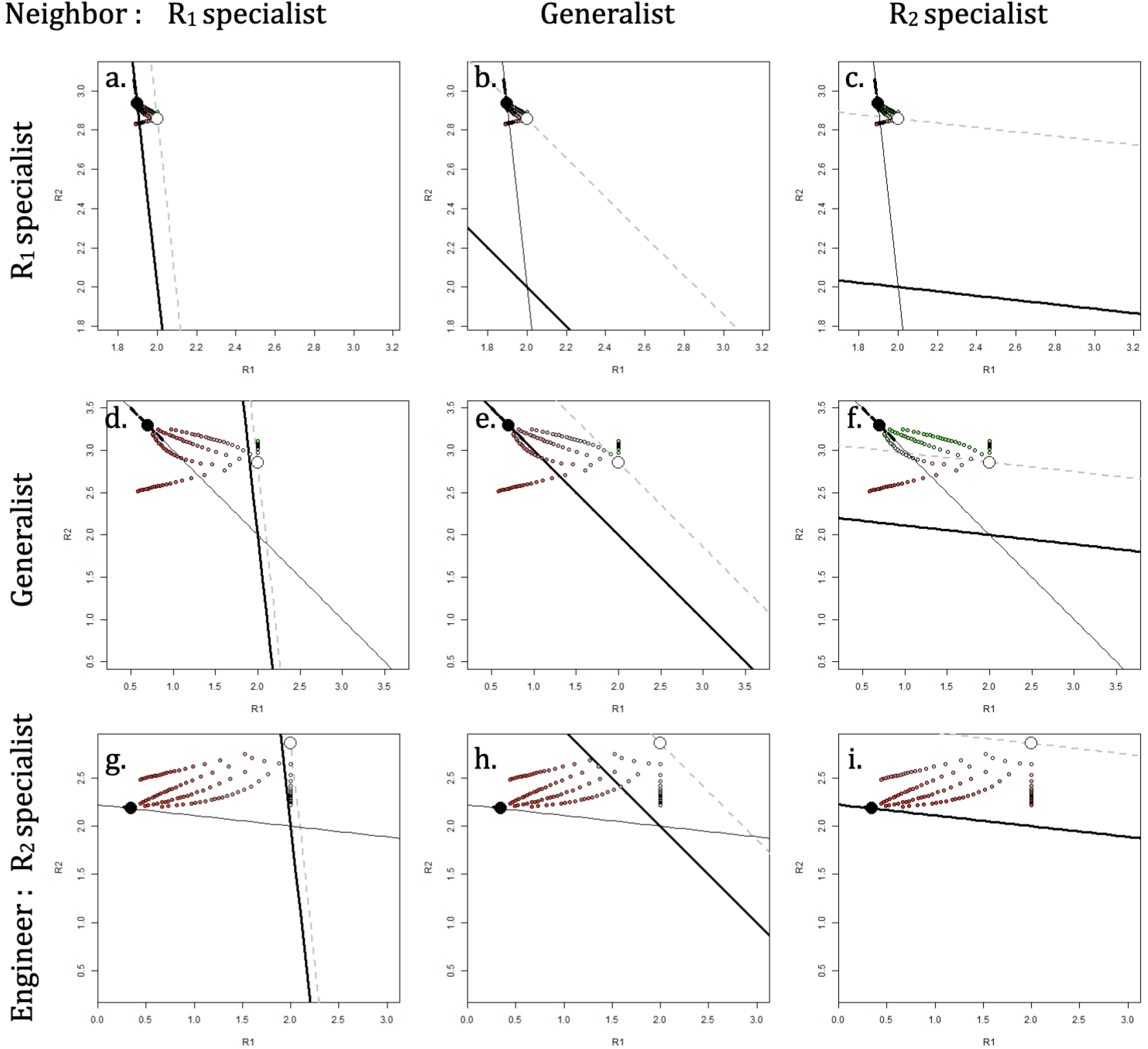
**Effect of an constructor accelerating the transformation of** *R*_1_ **into** *R*_2_. **This figure is symmetrical to Fig. C2 and C3. Here, the constructor has a positive effect on the richness of** *R*_2_, **the constructor has a positive effect as lon as the constructor is not specialised on** *R*_2_, **when** *R*_2_ **diffuses most and when the neighbour is specialised on** *R*_2_. *d*_0_ = 0.5, *α* = 5

## Appendix D: effect of the exchange rate and asymmetry in the farmer scenario

In this appendix, we explore more systematically how the effect of the constructor changes with the exchange rate and asymmetry. When the population of the constructor explodes, we can simply write the effect of the constructor on the neighbour Δ*r*_*N*_, and how that changes with the asymmetry in diffusion (Eq. D1) and the exchange rate *e* (Eq. D2) separately. When the population reaches an equilibrium, the expression of Δ*r*_*N*_ is more complex but a graphical exploration is possible (Fig. D1). When the population of the constructor explodes, the effect of asymmetry on Δ*r*_*N*_ is :

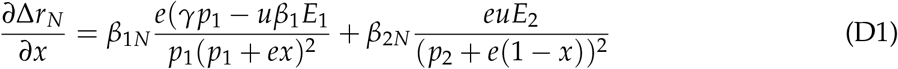

When condition 6 is fulfilled, *∂*Δ*r*_*N*_/*∂x* is always positive. This result i expected since *R*_1_ is the farmed resource : the more the asymmetry is biassed towards *R*_1_, the more the presence of the farmer is beneficial to the neighbour.

The effect of the exchange rate on Δ*r*_*N*_ is :

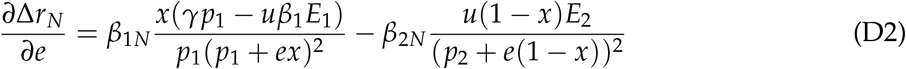

When the neighbour is a *R*_1_ specialist (*β*_1*N*_ = 1 and *β*_2*N*_ = 0) and that condition 6 is fulfilled, *∂*Δ*r*_*N*_/*∂e* is positive. On the contrary, when the neighbour is a *R*_2_ specialist (*β*_1*N*_ = 0 and *β*_2*N*_ = 1), *∂*Δ*r*_*N*_/*∂e* is negative. Due to the consumption of the non-farmed resource, the negative effect of the farmer increases as the exchange rate increases. For intermediate preferences, the effect of the exchange rate is not clear : increase in diffusion can be beneficial or detrimental depending on the gains of *R*_1_ relative to the losses in *R*_2_.

The constructor has a positive effect on the neighbour as soon as the neighbour is specialised on the farmed resource and when the exchange is sufficiently large. The effect of the exchange rate and the asymmetry on Δ*r*_*N*_ is represented on Fig. D1. For a given exchange rate *e* (vertical cut in Fig. D1), increase in *x* induces a shift from negative to positive effect (blue to beige zone), as described by Eq. D1. The result also holds for cases where the constructor reaches an equilibrium for which we don’t have an analytical expression (when *γ* ≤ 2). As for the effect of the exchange rate *e* (horizontal cut in Fig. D1), it is different depending on the level of niche construction *γ* and whether the neighbour is a generalist (panel a) or a specialist (panel b). On panel a, when *x* = 0.2 and the farmer has a strong effect (*γ* = 2.8), increasing the exchange rate induces a shift from negative to positive effect. On the contrary, when *x* = 0.6 and mild farming (*γ* = 2.2), increasing the exchange rate induces a shift from positive to negative.

**Figure D1:**
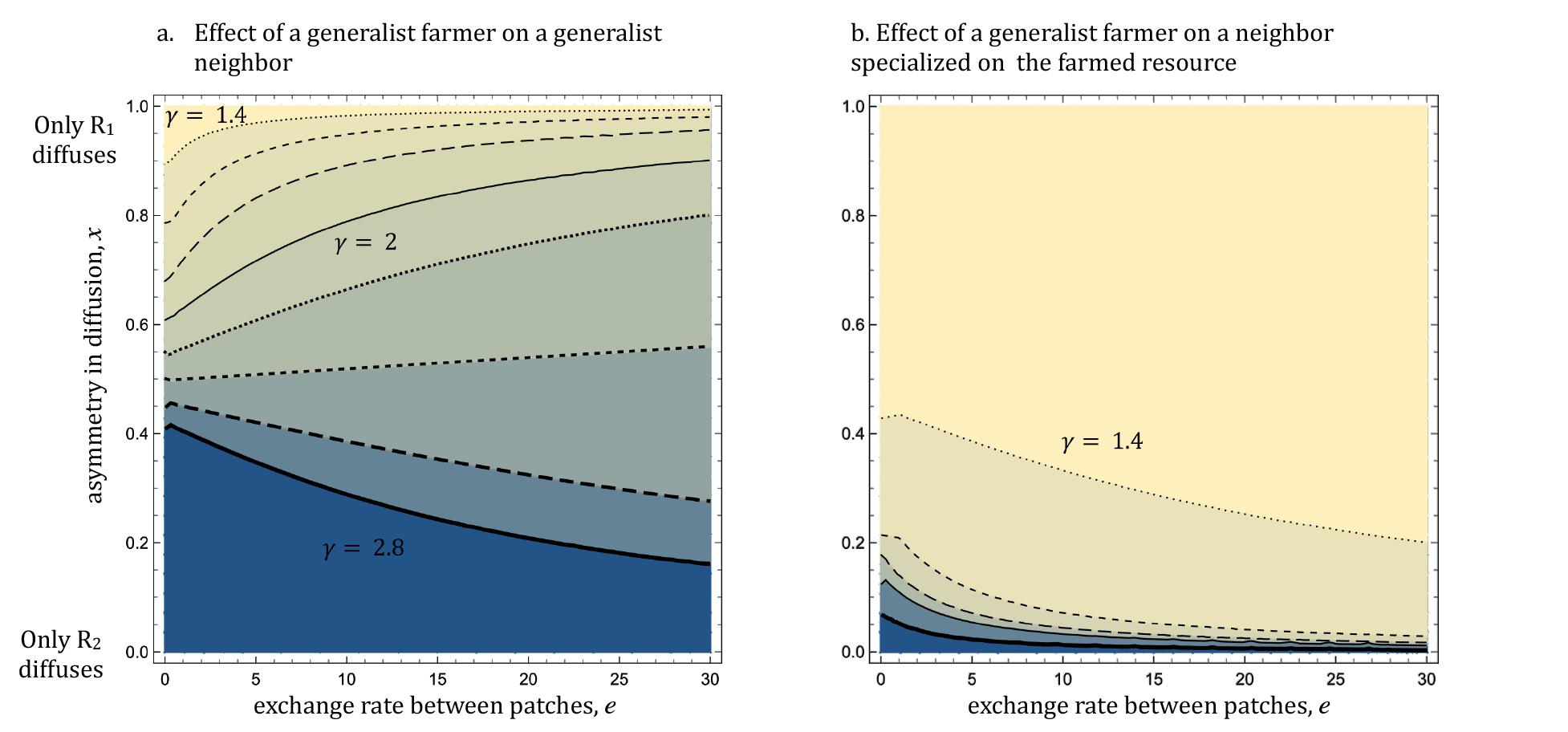
**The lines indicate situations where the farmer switches from having a facilitating effect (**Δ*r*_*N*_ > 0 **in beige) to having a negative effet (**Δ*r*_*N*_ < 0 **in blue), for different levels of farming intensities** *γ*. **a. Both the constructor and the neighbour are generalists (***β*_1_ = *β*_2_ = *β*_1*N*_ = *β*_2*N*_ = 0.5. **The exchange rate and asymmetry interact: at high levels of construction (***γ* < 2.6**), increase of diffusion and of asymmetry if beneficial for the neighbour ; At weak levels of construction (***γ* < 2.4**), increase of the exchange rate is detrimental for the neighbour. b. The constructor and the neighbour have different niches and facilitation is likelier. The constructor is a generalist and the neighbour a** *R*_1_ **specialist. Increase in diffusion or asymmetry is always beneficial**.

