## Appendices for "Diffusion of constructed resources and specialism promotes facilitation in spatial systems"

Supplementary files for: Ardichvili et al., 2024. Diffusion of constructed resources and specialism promotes facilitation in spatial systems. BioRXiv

#### Appendix A: feasibility analysis in the three scenario

From the model equations (1, 1b′ and (1b″-1e″), for every model the population equilibrium $C^*$ is the solution of a polynomial :

$$aC^{*3} + bC^{*2} + cC^* + d = 0 \tag{A1}$$

We use Descartes' rule to find a positive and real equilibrium.

In the three scenario,

$$\begin{aligned} d &= 0 \\ c &= \text{invasibility condition} * \text{positive term}(p_1, p_2, e, x) \\ b &= \text{complex expression} \\ a &= \text{simple expression} \end{aligned} \tag{A2}$$

$c$ is always positive when the invasibility condition is met.

Studying the sign of $a$ allows us to determine when an equilibrium is feasible.

In the three scenarios, $a$ reads $a_{conso}$, $a_\gamma$, and $a_\alpha$ for the consumer, farmer and transformer scenarios respectively.

$$
\begin{aligned}
a_{base} &= -p_C u^2 \beta_1 \beta_2 (p_1 + ex)(p_2 + e(1-x)) \\
a_\gamma &= (\gamma - p_C) u^2 \beta_1 \beta_2 (p_1 + ex)(p_2 + e(1-x)) \\
a_\alpha &= -p_C u \beta_2 (\alpha + u\beta_1)(p_1 + d_0 + ex)(p_2 + e(1-x))
\end{aligned}
\tag{A3}
$$

$a_{conso}$ is always negative, $a_\gamma$ is negative when $p_C > \gamma$, $a_\alpha$ is negative when $\alpha + u\beta_1 > 0$. The sign of $b$ does not inform us about the issue of the population dynamics since Descartes's rule cannot say whether there are - or 2 positive roots when there are two sign changes. The existence of 1, 2 or 0 equilibria (number in red) results in the following dynamics (in black):

| | $c > 0$<br>invasion is possible | | $c < 0$<br>nul eq. is stable | |
|---|---|---|---|---|
| | $b < 0$ | $b > 0$ | $b < 0$ | $b > 0$ |
| $a < 0$ | 1<br>$C^*$ | | 0<br>0 | 2 or 0<br>$0/C^*$ or 0 |
| $a > 0$ | 2 or 0<br>$C^*/\infty^+$ ou $\infty^+$ | 0<br>$\infty^+$ | 1<br>$0/\infty^+$ | |

$C^*$ indicates that the population stabilises at an equilibrium, 0 that extinction is locally stable, $\infty^+$ the the population grows exponentially. The / indicates situations of bi-stability where the dynamics depend on the initial abundance of the niche constructor.

### Appendix B: obtaining equations 5 and 6

#### *Equation 5*

At equilibrium, $R_{10} = E_1/p_1$, Eq. 1d = 0 and:

$$R^*_{1N} - R^*_{10} = \frac{ex}{p_1}(R^*_1 - R^*_{1N}) \tag{B1}$$

Assuming equilibrium, subtracting Eq. 1d to Eq. 1b yields

$$(1b) - (1d) = -u\beta_1 R^*_1 C^* + (R^*_{1N} - R^*_1)(p_1 + 2ex) = 0 \tag{B2}$$

$$(R^*_1 - R^*_{1N}) = \frac{-u\beta_1 R^*_1 C^*}{p_1 + 2ex} \tag{B3}$$

Subbing Eq.B3 into Eq. B1 yields Eq. 5.

#### *Equation 6*

In the second model, assuming equilibrium Eq. 1b′ and 1d can be re-arranged:

$$\begin{cases} (u\beta_1 C^* + p_1 + ex)R^*_1 - exR^*_{1N} = E_1 + \gamma C^* \\ -exR^*_1 + (p_1 + ex)R^*_{1N} = E_1 \end{cases} \tag{B4}$$

In matrix form, that yields

$$A \begin{pmatrix} R^*_1 \\ R^*_{1N} \end{pmatrix} = \begin{pmatrix} E_1 + \gamma C^* \\ E_1 \end{pmatrix} \tag{B5}$$

with

$$A = \begin{pmatrix} u\beta_1 C^* + p_1 + ex & -ex \\ -ex & p_1 + ex \end{pmatrix} \tag{B6}$$

And

$$(R_1^* - R_{1N}^*) = \frac{1}{det(A)}(\gamma p_1 - u\beta_1 E_1)C^* \tag{B7}$$

Where $det(A) = (p_1 + ex)C^*u\beta_1 + p_1(p_1 + 2ex)$ is positive, so from Eq. B7 and B1 yield condition 6 when the niche constructor reaches an equilibrium ($C^* > 0$).

When the niche constructor grows exponentially,

$$\lim_{C^* -> +\infty} R_1^* - R_{1N}^* = \frac{\gamma p_1 - u\beta_1 E_1}{u\beta_1(p_1 + ex)} \tag{B8}$$

and the same condition holds.

#### *Equation 7*

At equilibrium, $R_{10} = E_1/(p_1 + d_0)$, Eq. d" = 0 and:

$$R_{1N}^* - R_{10}^* = \frac{ex}{p_1 + d_0}(R_1^* - R_{1N}^*) \tag{B9}$$

Subtracting 1d" to 1b" :

$$(R_1^* - R_{1N}^*) = -\frac{(u\beta_1 + \alpha)C^* {R_1}^*}{d_0 + p_1 + 2ex} \tag{B10}$$

#### *Equation 8*

At equilibrium, $R_{20}^* = (E_2 + d_0 R_{10}^*)/(p_2)$.

Equation 1e" can be written :

$$\begin{aligned} \frac{dR_{2N}}{dt} &= E_2 + d_0 R_{1N} - d_0 {R_{10}}^* + d_0 {R_{10}}^* - p_2 R2N + e(1-x)(R_2 - R_{2N}) \\ &= p_2(R_{20} - R_{2N}) + d_0(R_{10}^* - R_{1N}) + e(1-x)(R_2 - R_{2N}) \end{aligned} \tag{B11}$$

At equilibrium, Eq. B11 $=0$ and we can write:

$$(R_{2N}^{*}-R_{20}^{*})=\frac{e(1-x)}{p_2}({R_2}^{*}-R_{2N}^{*})+\frac{d_0}{p_2}(R_{10}^{*}-R_{1N}^{*}) \tag{B12}$$

Subtracting 1e″ to 1c″ and assuming equilibrium, we obtain:

$$({R_2}^{*}-R_{2N}^{*})=\frac{1}{p_2+2e(1-x)}(-u\beta_2C^{*}R_2^{*}+\alpha C^{*}R_1^{*}+d_0(R_1^{*}-R_{1N}^{*})) \tag{B13}$$

After re-arranging the previous 4 equations, we get :

$$(R_{2N}^{*}-R_{20}^{*})=-Au\beta_2C^{*}R_2^{*}+A\alpha C^{*}R_1^{*}+\frac{d_0}{p_2}B(R_1^{*}-R_{1N}^{*}) \tag{B14}$$

with

$$\begin{aligned}A&=\frac{e(1-x)}{p_2(p_2+2e(1-x))}\\B&=\frac{e(1-x)}{p_2+2e(1-x)}+\frac{ex}{p_1+d_0}\end{aligned} \tag{B15}$$

$A$ and $B$ are both positive. The first term, $-Au\beta_2C^{*}R_2^{*}$ is the negative effect of the constructor due to the consumption of $R_2$. The second term, $A\alpha C^{*}R_1^{*}$ corresponds to the direct effect of the constructor on $R_2$ due to its niche construction activity; it is positive when the constructor is a stimulator and negative when the constructor is an inhibitor. The last term, $(d_0/p_2)B(R_1^{*}-R_{1N}^{*})$, is always negative when the niche constructor is a stimulator but can be positive when the niche constructor is an inhibitor. In other words, even an inhibitor can increase the concentration of $R_2$ in the empty patch.

### Appendix C: Effect of the niche constructor for different neighbour and constructor preferences

This appendix explores the effect of the niche constructor for different preferences $\beta_1$ and $\beta_{1N}$. Figure 3 in the main text shows several cases of facilitation. However, looking in more detail into several combinations of strategies highlight that cases of facilitation are rare.

First, facilitation is impossible when the constructor is specialised on the resource whose concentration increases due to niche construction ($R_1$ in the Farmer scenario or when the Transformer is an inhibitor; $R_2$ when the transformer is a Stimulator), which is in line with Eq. 6 and Eq.7. In other cases, facilitation is only possible when diffusion is sufficiently biassed towards the constructed resource. It's only when the constructor and the neighbour have sufficiently different niches that a broader range of exchange and asymmetry allow facilitation.

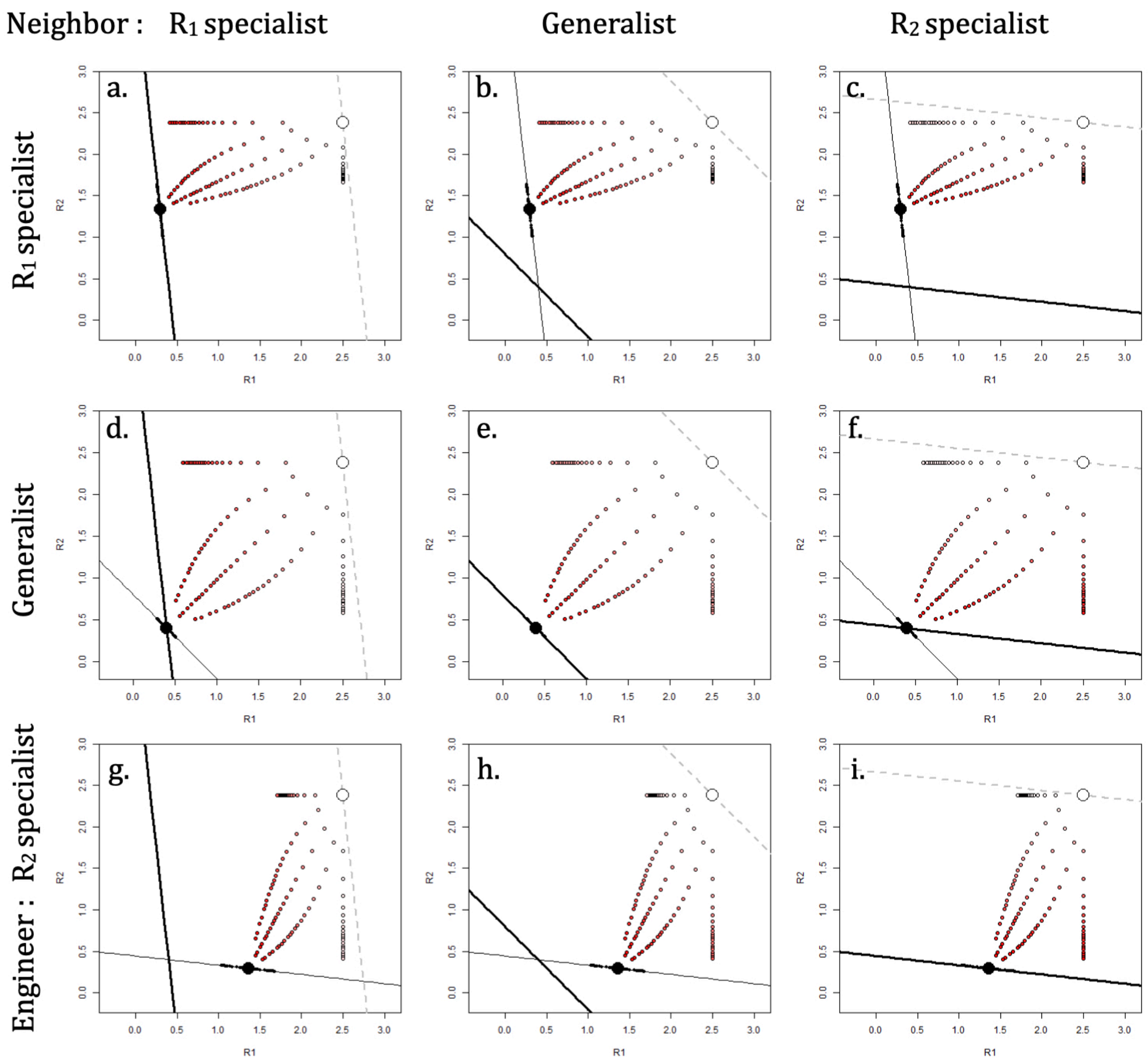


**Figure C1: Effect of the exchange rate and diffusion on the richness of the neighbour patch in the base model. $\Delta r_N$ is negative in red. As expected, whatever the strategies of the constructor or the neighbour, the effect is always negative. The colour scale is panel-dependent, darkest red corresponding to the worst case for the given $\beta_1$ and $\beta_{1N}$ combination.**

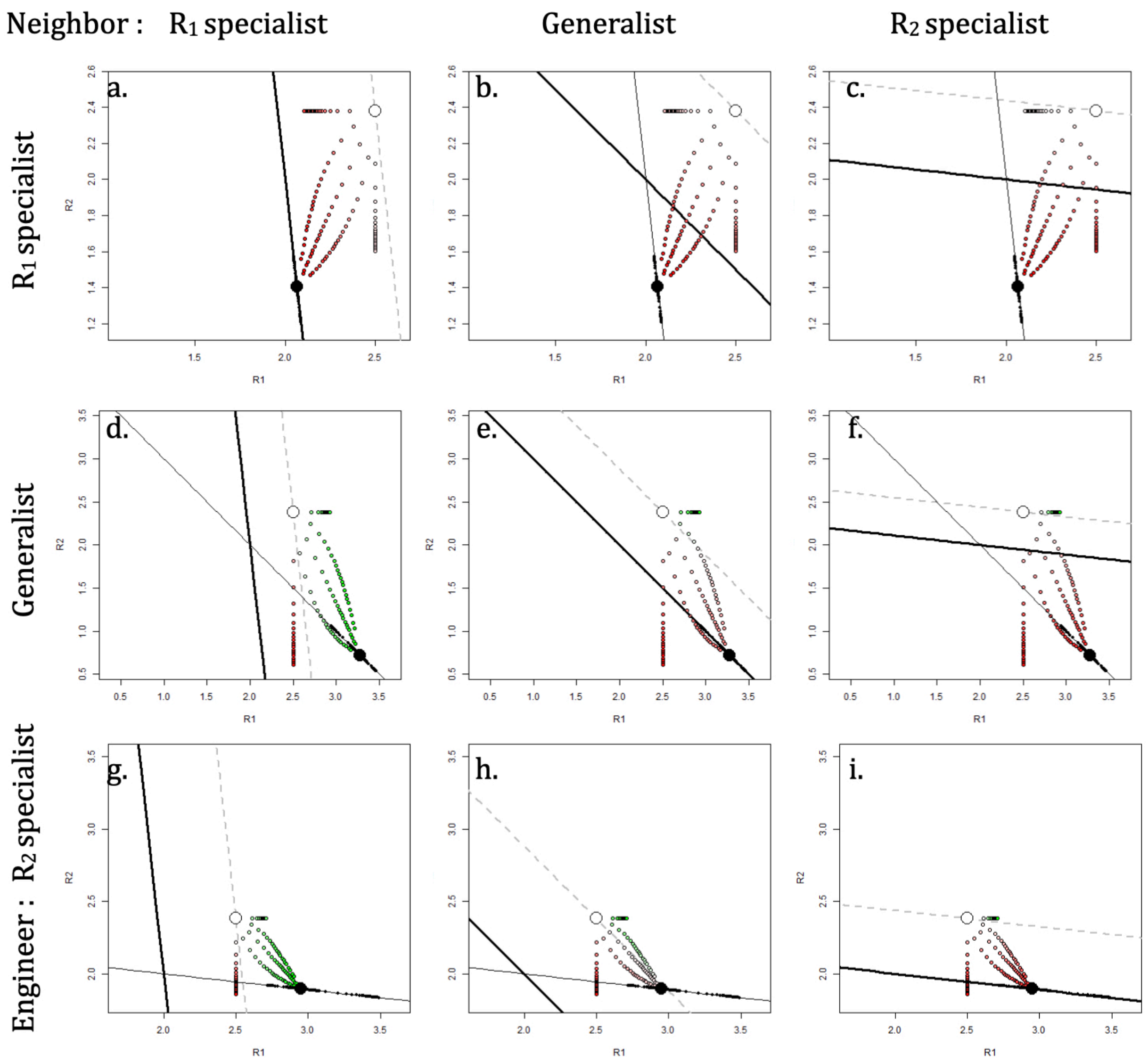


**Figure C2: Effect of a niche constructor farming resource 1. $\Delta r_N$ positive when the circle is green, negative when the circle is red. When the constructor is specialised on resource 1 (1st line), facilitation cannot happen (see condition 6). Facilitation is more likely when the neighbour has a strong preference for resource 1 (panels d and i). Level of facilitation :** $\gamma = 1.8$

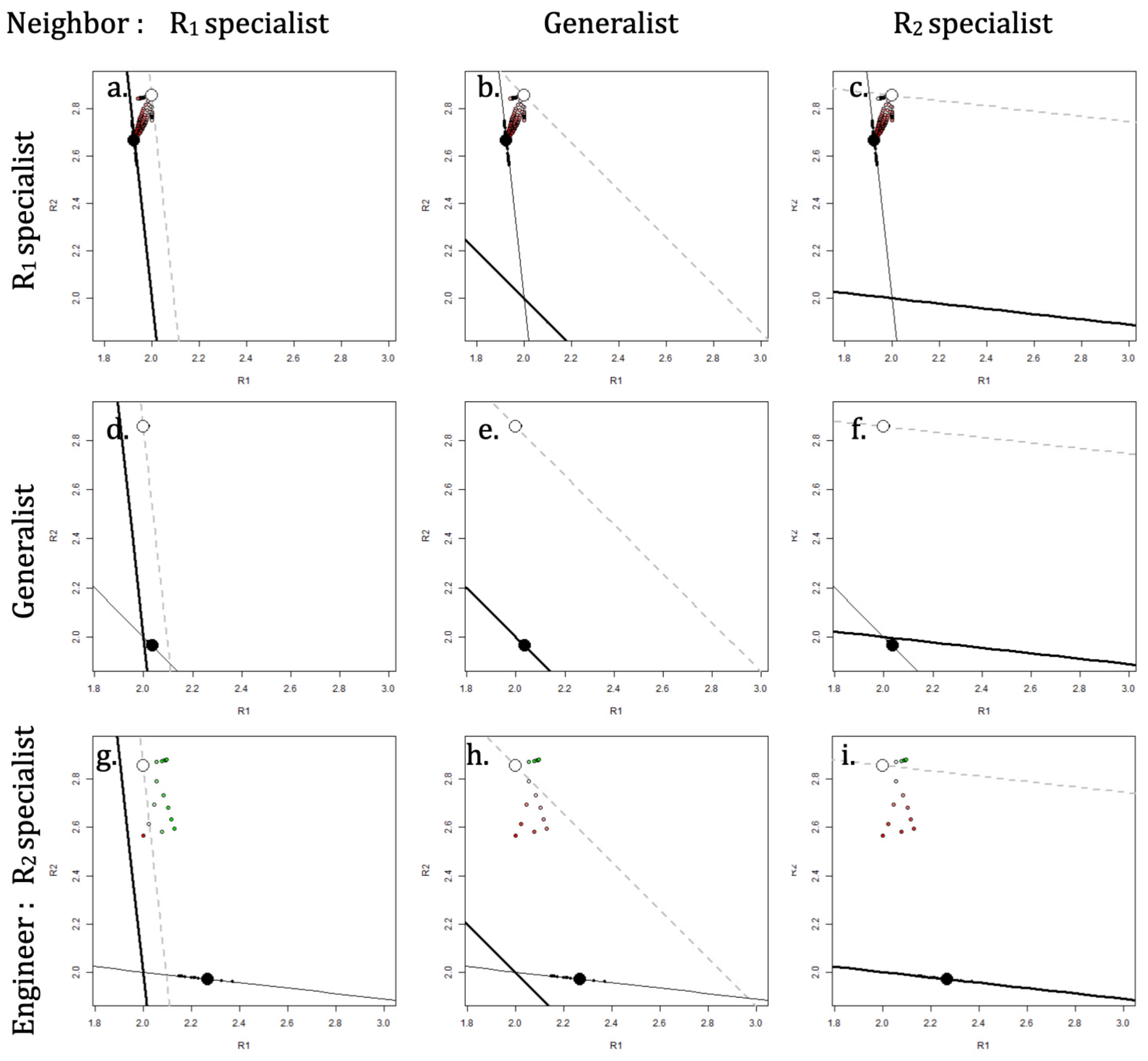


**Figure C3: Effect of an constructor slowing down the transformation of $R_1$ into $R_2$. As in the Farmer scenario, facilitation occurs only when the constructor is not a $R_1$ specialist. Additionally, the neighbour needs to be a $R_1$ specialist and only $R_1$ should diffuse.** $d_0 = 0.5$, $\alpha = -0.55$

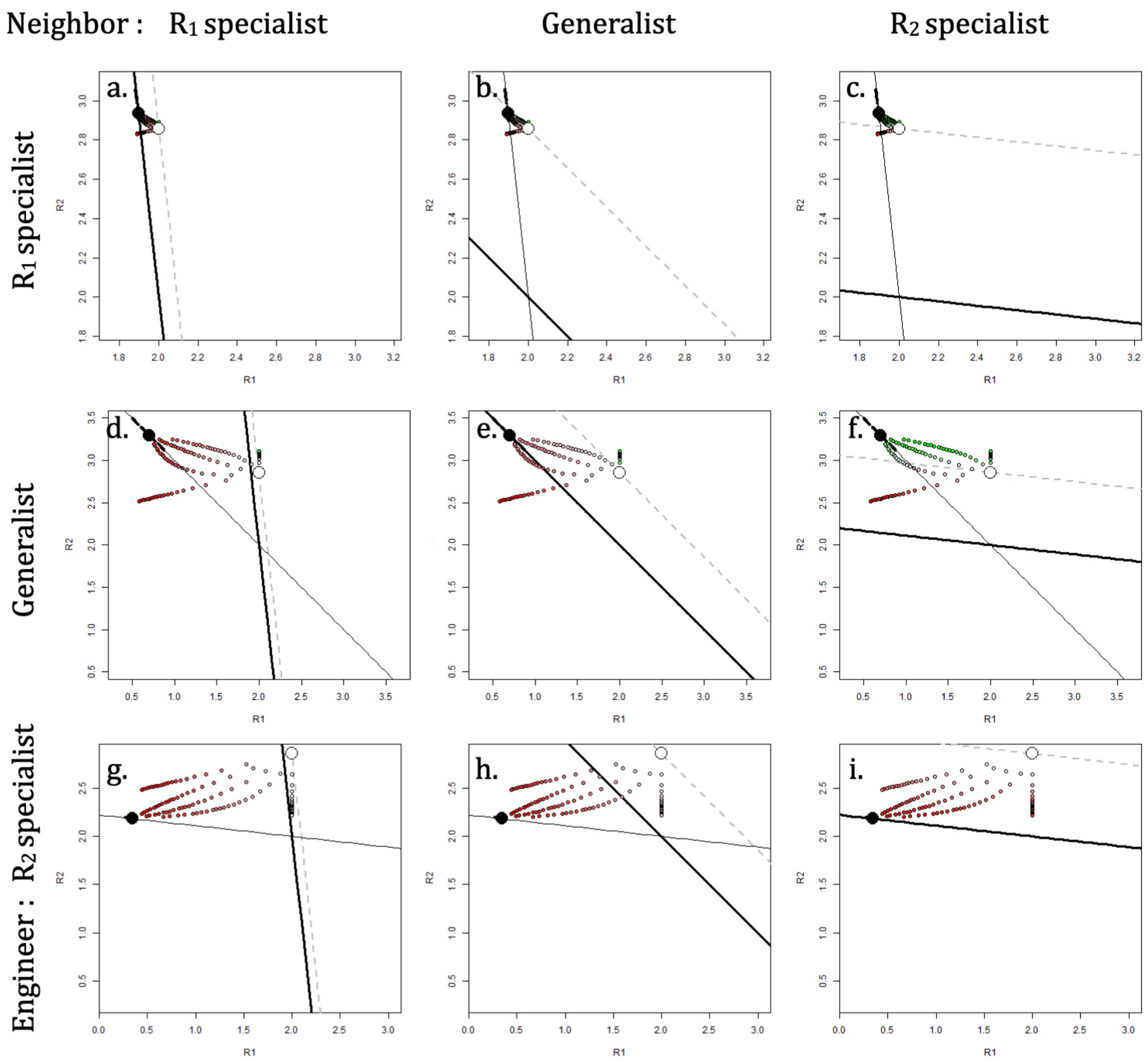


**Figure C4: Effect of an constructor accelerating the transformation of $R_1$ into $R_2$. This figure is symmetrical to Fig. C2 and C3. Here, the constructor has a positive effect on the richness of $R_2$, the constructor has a positive effect as lon as the constructor is not specialised on $R_2$, when $R_2$ diffuses most and when the neighbour is specialised on $R_2$.** $d_0 = 0.5$, $\alpha = 5$

### Appendix D: effect of the exchange rate and asymmetry in the farmer scenario

In this appendix, we explore more systematically how the effect of the constructor changes with the exchange rate and asymmetry. When the population of the constructor explodes, we can simply write the effect of the constructor on the neighbour $\Delta r_N$, and how that changes with the asymmetry in diffusion (Eq. D1) and the exchange rate $e$ (Eq. D2) separately. When the population reaches an equilibrium, the expression of $\Delta r_N$ is more complex but a graphical exploration is possible (Fig. D1). When the population of the constructor explodes, the effect of asymmetry on $\Delta r_N$ is :

$$\frac{\partial \Delta r_N}{\partial x} = \beta_{1N} \frac{e(\gamma p_1 - u\beta_1 E_1}{p_1(p_1 + ex)^2} + \beta_{2N} \frac{euE_2}{(p_2 + e(1-x))^2} \tag{D1}$$

When condition 6 is fulfilled, $\partial \Delta r_N / \partial x$ is always positive. This result i expected since $R_1$ is the farmed resource : the more the asymmetry is biassed towards $R_1$, the more the presence of the farmer is beneficial to the neighbour.

The effect of the exchange rate on $\Delta r_N$ is :

$$\frac{\partial \Delta r_N}{\partial e} = \beta_{1N} \frac{x(\gamma p_1 - u\beta_1 E_1)}{p_1(p_1 + ex)^2} - \beta_{2N} \frac{u(1-x)E_2}{(p_2 + e(1-x))^2} \tag{D2}$$

When the neighbour is a $R_1$ specialist ($\beta_{1N} = 1$ and $\beta_{2N} = 0$) and that condition 6 is fulfilled, $\partial \Delta r_N / \partial e$ is positive. On the contrary, when the neighbour is a $R_2$ specialist ($\beta_{1N} = 0$ and $\beta_{2N} = 1$), $\partial \Delta r_N / \partial e$ is negative. Due to the consumption of the non-farmed resource, the negative effect of the farmer increases as the exchange rate increases. For intermediate preferences, the effect of the exchange rate is not clear : increase in diffusion can be beneficial or detrimental depending on the gains of $R_1$ relative to the losses in $R_2$.

The constructor has a positive effect on the neighbour as soon as the neighbour is specialised on the farmed resource and when the exchange is sufficiently large. The effect of the exchange rate and the asymmetry on $\Delta r_N$ is represented on Fig. D1. For a given exchange rate $e$ (vertical cut in Fig. D1), increase in $x$ induces a shift from negative to positive effect (blue to beige zone), as described by Eq. D1. The result also holds for cases where the constructor reaches an equilibrium for which we don't have an analytical expression (when $\gamma \leq 2$). As for the effect of the exchange rate $e$ (horizontal cut in Fig. D1), it is different depending on the level of niche construction $\gamma$ and whether the neighbour is a generalist (panel a) or a specialist (panel b). On panel a, when $x = 0.2$ and the farmer has a strong effect ($\gamma = 2.8$), increasing the exchange rate induces a shift from negative to positive effect. On the contrary, when $x = 0.6$ and mild farming ($\gamma = 2.2$), increasing the exchange rate induces a shift from positive to negative.

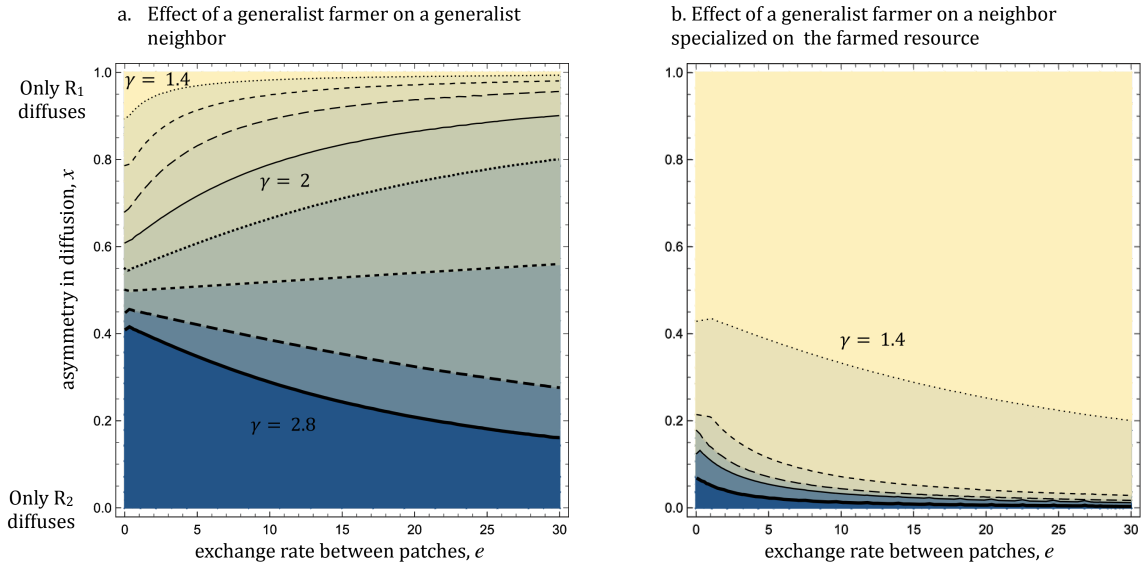


**Figure D1: The lines indicate situations where the farmer switches from having a facilitating effect ($\Delta r_N > 0$ in beige) to having a negative effet ($\Delta r_N < 0$ in blue), for different levels of farming intensities $\gamma$. a. Both the constructor and the neighbour are generalists ($\beta_1 = \beta_2 = \beta_{1N} = \beta_{2N} = 0.5$. The exchange rate and asymmetry interact: at high levels of construction ($\gamma < 2.6$), increase of diffusion and of asymmetry if beneficial for the neighbour ; At weak levels of construction ($\gamma < 2.4$), increase of the exchange rate is detrimental for the neighbour. b. The constructor and the neighbour have different niches and facilitation is likelier. The constructor is a generalist and the neighbour a $R_1$ specialist. Increase in diffusion or asymmetry is always beneficial.**
